# NOTCH1-specific phosphorylation of S1970 by Casein Kinase 1 is required for NOTCH1 transcriptional competence and signaling activity *in vivo*

**DOI:** 10.64898/2026.05.05.722849

**Authors:** Fabio Turetti, Lorena Agostini Maia, Hana Hajšmanová, Jakub Sławski, Wim Dehaen, Marek Dokoupil, Giovanna Collu, Tomáš Gybeĺ, Kamil Paruch, Emma R. Andersson, Vítězslav Bryja, Pavla Perlikova, Konstantinos Tripsianes, Jakub Harnoš, Jan Mašek

## Abstract

The Notch and Wnt/β-catenin signaling pathways are essential regulators for cell-fate decisions, cellular patterning, and tissue homeostasis. Multiple studies point to their orchestrated role during development, but the molecular mechanism of the protein-protein crosstalk is largely unknown. Here, after screening effects of Wnt/β-catenin component loss on NOTCH1 protein, we identify Casein Kinase 1α (CK1α) as a positive regulator of NOTCH1 activity *in vitro* and *in vivo*. We demonstrate that CK1α associates with NOTCH1 and that its kinase activity is required to sustain Notch-driven transcription. Using UltraID proximity-assay, we revealed that CK1α is required for the NOTCH1 interactivity with transporter proteins, and MAML1 both prior and after ligand-induced activation. Combining structural modelling, NMR, and mass spectrometry, we identified Serine 1970 (S1970) as a previously unreported residue within the Notch1 Intracellular Domain (N1ICD) essential for its signaling competence. Our modeling predicts that the phosphorylation of S1970 facilitates an intra-domain conformational switch with R1937 and R1962 residues altering the assembly of the N1ICD-MAML1-RBPJk transcriptional complex. Finally, we demonstrate the biological significance of N1ICD S1970 *in vivo* using *Xenopus laevis* axis-duplication rescue assay. Our results establish CK1α as a key positive mediator of the Notch receptor transcriptional activity.

## INTRODUCTION

The coordination of the Notch and Wnt/β-catenin signaling pathway, frequently referred to as “Wntch” crosstalk (Hayward et al., 2008; Turetti et al., 2026), is a fundamental requirement for precise cellular patterning (Couso & Arias, 1994), tissue homeostasis (Tian et al., 2015), and cell-fate decisions (Kwon et al., 2009). Despite their ubiquity, the molecular dependencies between the Wnt and Notch pathways remain characterized by contradicting observations (Turetti et al., 2026), appearing synergistic in some cellular contexts and antagonistic in others (Collu et al., 2014). The lack of our insight into precise molecular mechanism by which components of the two pathways physically interact to fine-tune the cell differentiation prevents us from testing the importance of Wntch *in vivo*.

The Wnt/β-catenin signaling activity is primarily dictated by the amount of β-catenin stability, of which is tightly controlled by the cytoplasmic “destruction complex”—a multi-protein scaffold comprising Axin, Adenomatous polyposis coli (APC), β-transducin repeat-containing protein (β-TrCP), Glycogen synthase kinase-3 beta (GSK3β), and Casein Kinase 1α (CK1α) (Clevers et al., 2014; Ranes et al., 2021). In the absence of Wnt ligands, the destruction complex targets β-catenin for phosphorylation and subsequent proteasomal degradation. Upon Wnt ligand binding to the Frizzled receptors and co-receptors such as low-density lipoprotein receptor-related protein 5 and 6 (Lrp5/6), the destruction complex is relocated on the plasma membrane by Dishevelled proteins (Dvls), allowing β-catenin to accumulate, translocate to the nucleus, and subsequently activate transcription *via* TCF/LEF transcription factors (Clevers et al., 2014; Gammons & Bienz, 2018).

Conversely, Notch signaling is triggered by juxtacrine ligand binding from a neighboring, “sending cell”. The pulling force generated by the ligand endocytosis induces sequential proteolytic cleavages of the Notch receptor. The final (S3) cleavage, mediated by γ-secretase (De Strooper et al., 1999; Schroeter et al., 1998), releases the Notch Intracellular Domain (NICD) from the plasma membrane in the “receiving cell“. The NICD then translocate to the nucleus to form the ternary transcriptional complex with the DNA-binding protein RBPJk (also known as CSL) and the co-activator Mastermind-like (MAML), triggering expression of Notch target genes, such as *HES* (Jarriault et al., 1995) and *HEY* families (Bray & Bigas, 2025; Leimeister et al., 2000).

Unlike Wnt/β-catenin, the Notch pathway lacks signal amplification; each ligand can induce release of only one NICD molecule. Consequently, posttranslational processing and stability-regulators of NICD are among the key determinants of the Notch signaling output, which, when deregulated, drives genetic disorders or cancer (Bray & Bigas, 2025; Mašek & Andersson, 2017). Among the central regulators of the NICD stability is proteasomal degradation governed by SCF-FBX7 (Hoeck et al., 2010). The activity of SCF-FBX7 though, was shown to be regulated by NICD phosphorylation by kinases, such as GSK3β (Foltz et al., 2002a; Han et al., 2012), CDK8 (Fryer et al., 2004), or CDK1/2 (Carrieri et al., 2019). In addition to affecting protein stability, phosphorylation of the NICD can affect also the assembly of the transcriptional complex itself. CK2-mediated phosphorylation has been shown to reduce the affinity of N1ICD for its co-factors MAML1 and RBPJk, thereby attenuating the Notch-response (Ranganathan et al., 2011).

In this study, we identify Casein Kinase 1α (CK1α) as a novel positive regulator of Notch signaling both *in vitro* and *in vivo*. We demonstrate that CK1α associates with NOTCH1 and is required for its transcriptional activity independently of the Wnt stimulation. We further identified a previously unreported residue, Serine 1970 (S1970) in N1ICD ANK3 repeat to be phosphorylated by CK1 and confirmed its importance for N1ICD activity *in vivo*. Finally, using a combination of proximity labelling, AlphaFold, and computational modelling, we propose that the phosphorylated S1970 in ANK is required for interaction with R1937 ANK2 domain, necessary for the N1ICD-MAML1-RBPJk transcriptional complex assembly.

## RESULTS

### Several core Wnt/β-catenin components affect protein levels of active, S3-cleaved NOTCH1

Various cell lines and approaches were used to address the interaction between individual NOTCH receptors and individual components of the Wnt/β-catenin pathway, but no side-by-side comparison has been performed so far (Turetti et al., 2026). To probe the extent of effect of individual WNT components on NOTCH1 receptor biology comparably, we performed a small-scale screen in HEK TREx cells devoid of i) either individual upstream regulators of the β-catenin Destruction complex like *DVL1/2/3*^-/-^ (Paclíková et al., 2017), *LRP5/6*^-/-^, or *RNF43/ZNRF3*^-/-^ (Radaszkiewicz et al., 2020) or ii) proteins of the Destruction complex itself (*AXIN1/2*^-/-^, *GSK-3β*^-/-^ or *CK1α*^-/-^ (Gybeľ et al., 2024)) (**Fig. 1A**).

**Figure 1:**
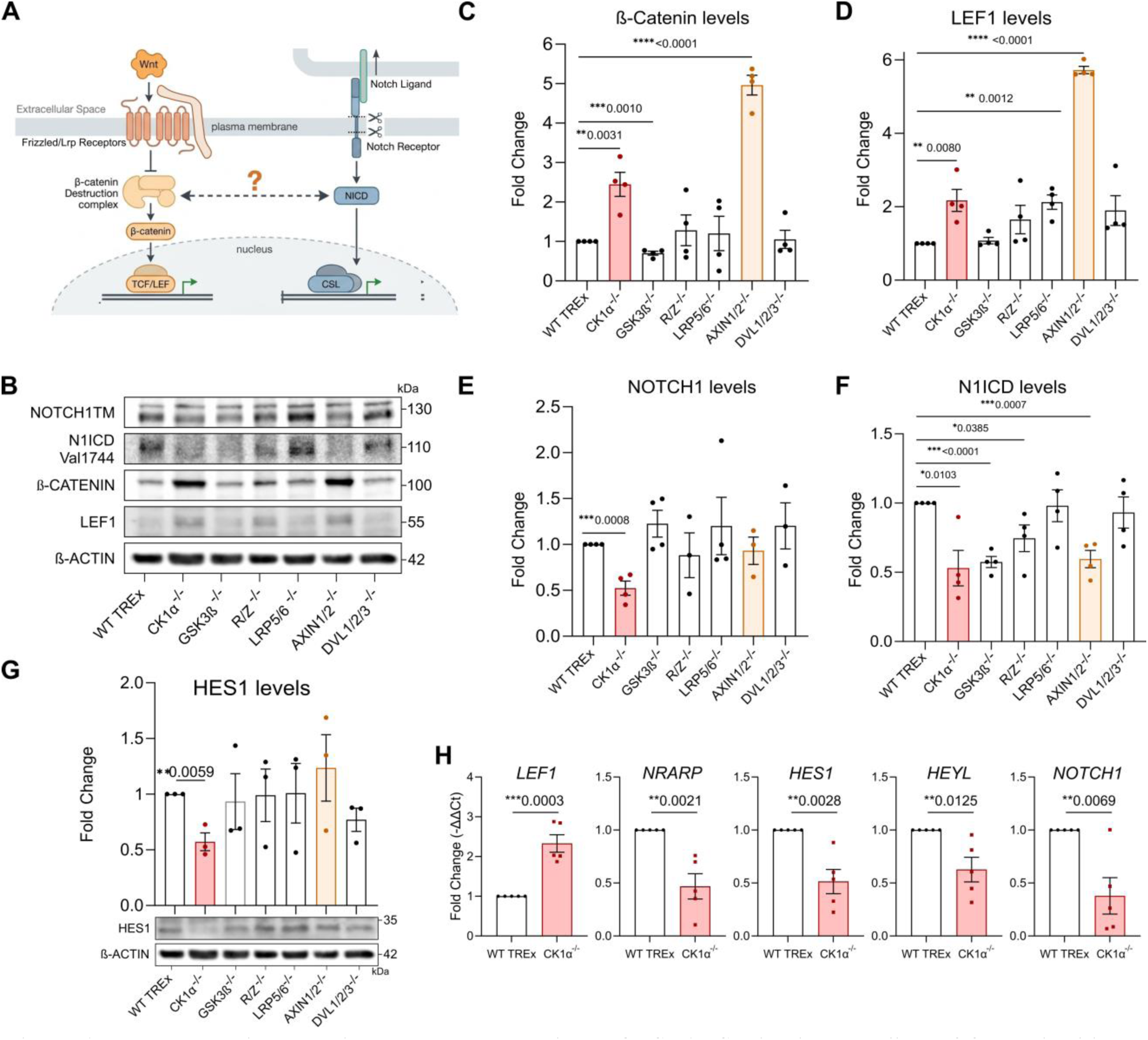
Wnt Destruction complex components regulate NOTCH1 ICD levels, regardless of β-catenin-driven transcriptional activity. (**A**) Scheme of the tested Wnt x Notch interaction. (**B-F**) Western blot of the transmembrane NOTCH1, cleaved NOTCH1 ICD (Val1744), total β-CATENIN, and LEF1 in the control HEK TREx and a panel of WNT KO cells (A), with quantifications (E-F), β-ACTIN was used for normalization. (**G**) Western blot and quantification of HES1 in the control HEK TREx and a panel of WNT KO cells (**H**) qPCR of Wnt (LEF1) and Notch (HES1, HEYL, NRARP, NOTCH1) target genes in the control HEK TREx and CK1α^-/-^ cells. R/Z, RNF43/ZNRF3; Each dot represents a biological replicate; Unpaired-T test (* P ≤ 0.05; ** P ≤ 0.01; *** P ≤ 0.001; **** P ≤ 0.0001, ns = not significant); Bars represent SEM.

In steady-state (i.e., not experimentally induced Wnt or Notch activation), the loss of components upstream of the destruction complex (DVL1/2/3, RNF43, ZNRF3, and LRP5/6) had little, or no effect on the β-CATENIN and its Wnt downstream target LEF1, a well-established readout of pathway activity (Doumpas et al., 2018) (**Fig. 1B**). On the other hand, deletion of the core components of the Destruction complex CK1α (Liu et al., 2002), and AXIN1/2 (Behrens et al., 1998; Hart et al., 1998; Peifer et al., 1994) lead to a respective 2.45-, and 4.96-fold increase of β-CATENIN protein levels, and corresponding 2.17-, and 5.72-fold enrichment of LEF1 (**Fig. 1C, D**). These results suggest that AXIN1/2 and CK1α are involved in β-CATENIN protein level maintenance, in agreement with previous results (Clevers et al., 2014; Ranes et al., 2021).

We next assessed the levels of endogenous NOTCH1TM (TM-transmembrane), and activated—S3 cleaved—N1ICD, using Val1754-specific antibody (**Fig. 1B**), revealing a significant NOTCH1TM and N1ICD decrease in *CK1α*^-/-^ (0.52-, 0.53-fold of the control, respectively), and respective N1ICD-selective decrease to 0.57-, 0.59-fold and 0.75-fold of the control in *GSK-3β*^-/-^, *AXIN1/2*^-/-^ and *R/Z^-/-^* cells (**Fig. 1E, F**). The absence of change in NOTCH1TM in *AXIN1/2*^-/-^ implies the observed effects are not a result of aberrant β-CATENIN-driven transcription (**Fig. 1C, D**), which is in line with observations from *Drosophila* showing Armadillo does not affect the total levels of Notch receptor (Muñoz-Descalzo et al., 2011). Strikingly, *CK1α*^-/-^ cells manifested a significant decrease of both NOTCH1 TM and N1ICD, as well as Notch target HES1; however, HES1 levels were unchanged in *GSK-3β*^-/-^, *AXIN1/2*^-/-^ cells (**Fig. 1G**). The changes in protein levels were accompanied by corresponding changes in mRNA; the Wnt target gene *LEF1* was significantly upregulated, while Notch target genes *NRARP*, *HES1, HEYL*, and *NOTCH1 (Castel et al., 2013; Chen et al., 2017; Wang et al., 2011)* were downregulated in *CK1α^-/-^*cells, when compared to the WT controls (**Fig. 1I**). This suggest a wider CK1α-specific role in NOTCH1 biology, which we investigated further.

### CK1α kinase activity stimulates N1ICD-driven transcription independently of Wnt activation

To investigate further the effect of CK1α on both N1ICD and Wnt-driven transcription, we performed Super TOPFlash (Veeman et al., 2003) (Wnt reporter) and 12xCSL (Kato et al., 1997) (Notch reporter) luciferase-based reporter assay in WT TREx, *CK1α*^-/-^. The *AXIN1/2*^-/-^ cell line served as control with “high” β-catenin signaling. In accord with β-CATENIN and LEF1 accumulation (**Fig. 1B-D**), loss of CK1α and AXIN1/2 resulted in hyperactivation of the Wnt reporter, showing a high amplitude range of signaling activation, of 169.70 and 858.70-fold of the control, respectively **(Fig. 2A, S1A**).

**Figure 2:**
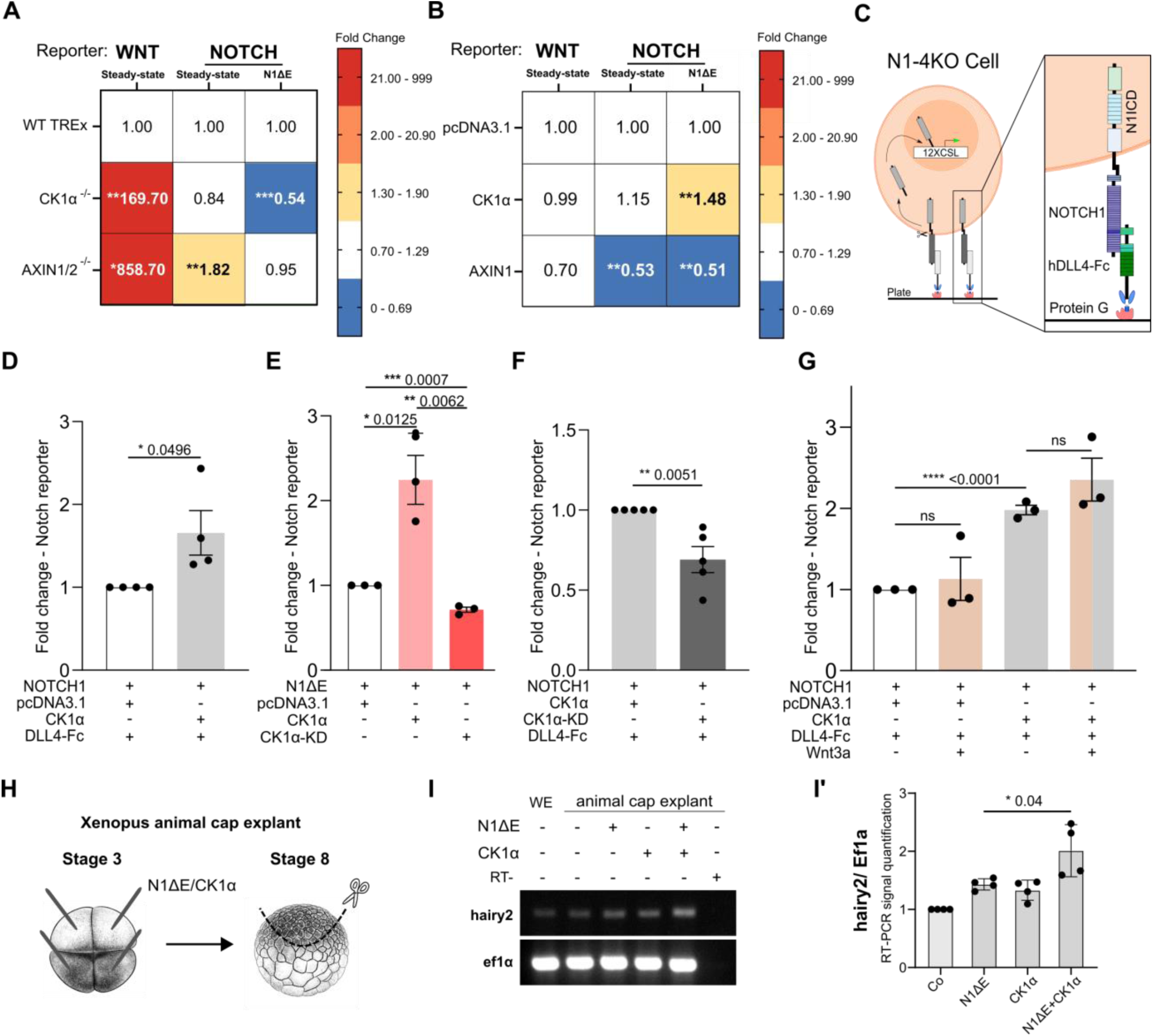
CK1α associates with the Notch to sustain signaling and regulate NICD activity (A-B) Heatmap visualization of WNT and NOTCH luciferase reporter assays, in steady state or after N1ΔE overexpression in CK1α^-/-^, AXIN1/2^-/-^ and control HEK TREx cells (A), or in WT HEK 293T following N1ΔE co-overexpression with selected WNT components (B). **(C)** Scheme of the Fc-DLL4 coating experiment. **(D-F)** Luciferase Notch reporter assay in N1-4KO cells on Fc-DLL4 coated plates transfected with NOTCH1 and pcDNA3.1 or CK1α (D), or CK1α-KD (E), or in HEK293T cells, transfected with NIΔE and either pcDNA3.1, CK1α, or CK1α-KD (E). **(G)** The **s**ame assay as in D treated with recombinant Wnt3a (150ng/ul) or PBS as control. (**H-I’**) Design of the Xenopus laevis animal cap explant experiment (H). Representative RT-PCR analysis of the hairy2 (Hes4 homolog in X. laevis) in animal cap explants. Ef1α was used as a loading control, and RT− lane served as a negative control (I) Densitometric quantification of RT-PCR bands intensity (hairy2 normalized to Ef1α (I’). AU, arbitrary units; Data are presented as mean with SD; Each dot represents a biological replicate; Unpaired-T test (* P ≤ 0.05; ** P ≤ 0.01; *** P ≤ 0.001, *** P ≤ 0.0001, ns = not significant); Bars represent SEM.

In uninduced state, the absence of CK1α had no significant effect on Notch reporter signal, while the steady-state activity of Notch reporter activity in *AXIN1/2^-/-^* cells was increased (1.82-fold of the control levels) (**Fig. 2A, S1B**). However, when we induced the Notch signaling by a constitutively active form of mN1ICD (N1ΔE), we observed a significantly decreased Notch reporter activity in *CK1α*^-/-^ compared to control TREx cells (0.54-fold of the control) (**Fig. 2A, S1B’**). In contrast, we detected no difference in Notch activity between the N1ΔE-transfected AXIN*1/2*^-/-^ and control cells (**Fig 2A, S1B’**), suggesting CK1α acts as a positive regulator of N1ICD, while AXIN1/2 as a negative regulator of NOTCH1.

To further test this possibility, we overexpressed CK1α or AXIN1, with or without N1ΔE, in HEK293T cells measuring the Notch or a Wnt reporter activity 24h later. The CK1α overexpression had no significant effect on Wnt (Borgal et al., 2014) nor on Notch-driven transcription (**Fig. 2B, S1D**), implicating the system is already saturated in steady-state. However, after activation of the Notch signaling with N1ΔE, CK1α enhanced Notch reporter activity by 1.48-fold (**Fig. 2B, S1C’**). AXIN1 overexpression resulted in a significant reduction of Notch reporter activity (0.53-fold of the control) under non-induced conditions, and 0.51-fold of the control after N1ΔE-mediated Notch activation (**Fig. 2B, S1C-C’**), implying it acts on NOTCH1 activity either indirectly or prior to NOTCH1 S3 cleavage.

Seeking further validation in a system with more physiological activation of the full length NOTCH1, we employed the recombinant Notch ligand coating assay (**Fig. 2C**) with Fc-Delta like 4 (DLL4) – the most potent activator of NOTCH1 (Kakuda et al., 2020). To mitigate potential effects on other Notch receptors, we utilized HEK293T cell line deficient in all four NOTCH receptors (N1-4KO), transfected with NOTCH1 alone, or with CK1α and assessed Notch reporter activity following cultivation on control (only Fc) and Fc-DLL4-coated plates. Co-expression of CK1α enhanced Notch reporter activity by 1.66-fold (**Fig. 2D**), which was consistent with Fig. 2B.

Next, we tested whether CK1α affects NOTCH1 signaling via its kinase activity. We co-transfected HEK293T cells, with N1ΔE and wild-type CK1α or a kinase-dead (KD) CK1α variant (Jiang et al., 2018). The KD-CK1α significantly reduced the Notch reporter activity (0.71-fold of the control), acting as dominant-negative (**Fig. 2E**). We observed similar effects also in N1-4KO cells, seeded on a Fc-DLL4-coated plate, with NOTCH1 and either wild-type CK1α or KD-CK1α (**Fig. 2F**), suggesting that CK1α-mediated phosphorylation is involved in N1ICD signaling.

Finally, to test if the CK1α effect on NOTCH1 is dependent or independent of Wnt activity, we performed a Fc-DLL4 coating assay in NOTCH1-transfected N1-4KO cells treated with recombinant Wnt3a or PBS. Neither the control, Fc-DLL4-NOTCH1, nor CK1α-enhanced activation of Notch reporter activity were significantly affected by presence of Wnt3a ligand (**Fig. 2G**). To see if the effect of CK1α on NICD is relevant also *in vivo*, we injected N1ΔE alone or with CK1α into blastomeres of a 4-cell stage *Xenopus laevis* embryo, let it develop until the blastula stage (NF stage 8), when we dissected and cultured the animal caps until stage 10.5 (**Fig. 2H**). We next analyzed the Notch target gene *Hes4* (*hairy2* in *Xenopus*) (Jarriault et al., 1995), and in line with results from HEK cells, co-injection of N1ΔE with wild-type CK1α led to significant increase in *hairy2* expression (**Fig. 2I, I’**). These data suggest that, independently of the established inhibitory role in Wnt signaling (Amit et al., 2002; C. Liu et al., 2002), CK1α is also required for N1ICD signaling activity.

### Proximity labeling reveals that CK1α degradation impairs NOTCH1 interaction with core signaling and trafficking proteins

Several kinases, including GSK3β (Han et al., 2012; Kim et al., 2009), and CDK1/2 (Carrieri et al., 2019) were shown to phosphorylate NICD, targeting it for proteasomal degradation, thus acting as its negative regulators. On contrary, co-transfecting HEK293T cells with N1ΔE and CK1α promoted N1ICD protein levels by 1.68-fold of the control (**Fig. 3A, A**’), matching the N1ICD decrease in CK1α^-/-^ cells (**Fig. 1F**). Importantly, over-expression of AXIN1 with N1ΔE did not affect N1ICD protein levels, reinforcing the notion of AXIN1 effects described in Figs. 1 and 2 are indirect (**Fig. 3A, A’**). To investigate whether the CK1α-dependent increase in N1ICD stability is due to increase N1ICD protein stability, we treated WT and CK1α^-/-^ cells, transfected with low amount of N1ΔE, with the inhibitor of translation, cycloheximide. We observed a ∼50% drop of N1ICD protein levels six hours after cycloheximide treatment in both conditions suggesting there is no difference in the N1ICD protein stability in *CK1α^-/-^* and control cells (**Fig. 3B**). This indicates that CK1α is involved in *NOTCH1* expression rather than protein stability per se.

**Figure 3:**
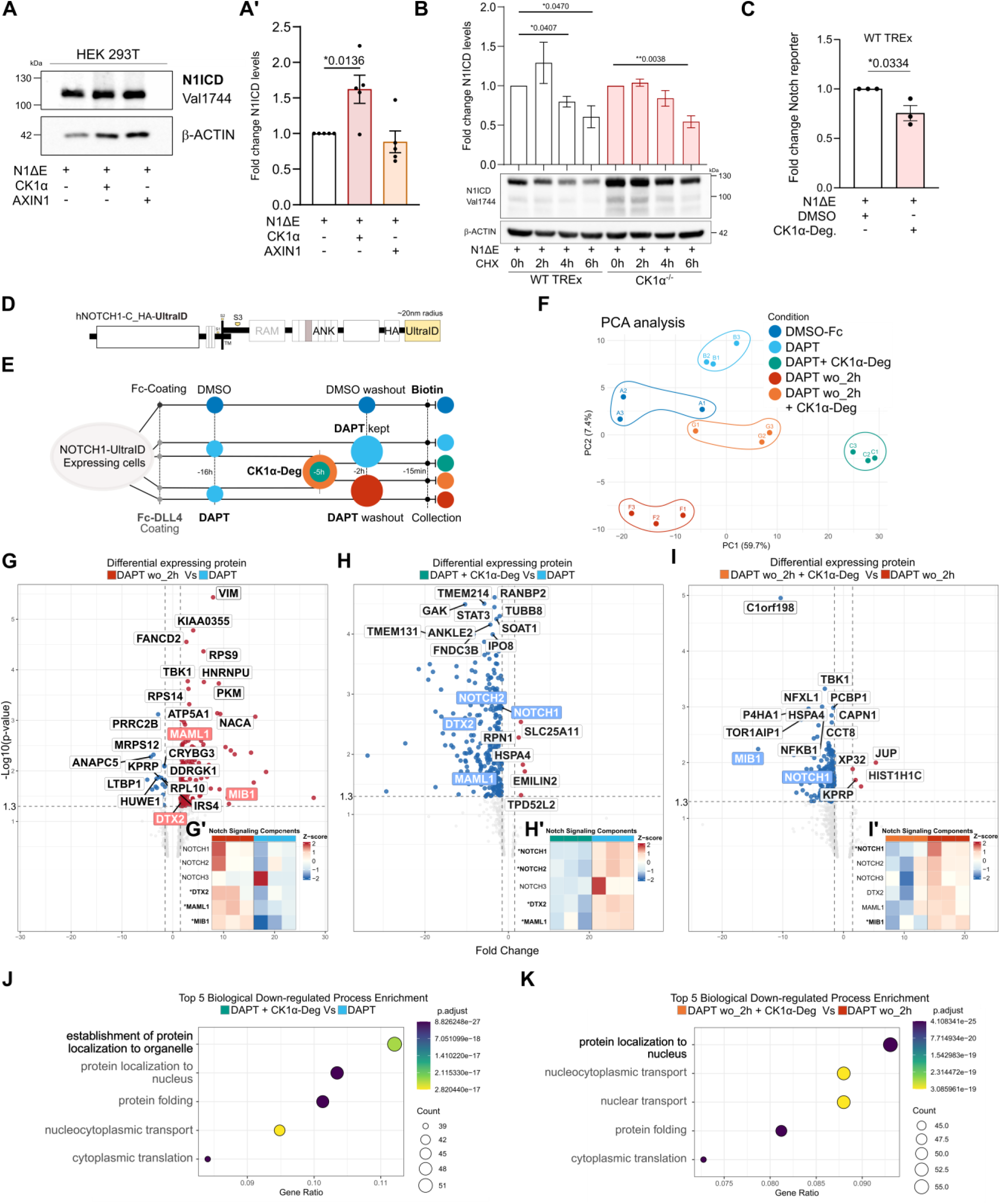
CK1α degradation impairs Notch trafficking proximity. (**A**) Western blot of the cleaved NOTCH1 ICD (Val1744), in WT HEK 293T transfected with N1ΔE and either CK1α or AXIN1 with (**A’**) protein quantification. β-ACTIN was used for normalization. (**B**) Western blot and protein quantification of cleaved NOTCH1 ICD (Val1744), in HEK TREx and CK1α^-/-^ cells transfected with N1ΔE and treated with either DMSO (6h) or Cycloheximide (CHX) (100µg/mL) for 2,4 or 6h. (N=3), β-ACTIN was used for normalization. (**C**) Luciferase Notch reporter assay in HEK TREx cells transfected with N1ΔE and treated with either DMSO or CK1α degrader (0.01µM). (**D**) Scheme representing hNOTCH1-UltraID construct expressed by Flp-in HEK 293 cells. (**E**) Experimental design of the hNOTCH1-UltraID proximity-dependent biotinylation assay. (**F**) Principal Component Analysis (PCA) of all the conditions. (**G**) Volcano plot visualization of the DEP in DAPT wo_2h Vs DAPT groups, and (**G’**) heatmap visualization of the Notch signaling components identified. (**H**) Volcano plot visualization of the DEP in DAPT + CK1α-Deg Vs DAPT groups, and (**H’**) heatmap visualization of the Notch signaling components identified. (**I**) Volcano plot visualization of the DEP in DAPT wo_2h + CK1α-Deg Vs DAPT wo_2h, and (**I’**) heatmap visualization of the Notch signaling components identified. (**J**) Dot plot visualization of the top 5 down-regulated Biological Process in DAPT + CK1α-Deg Vs DAPT groups. (**K**) Dot plot visualization of the top 5 down-regulated Biological Process in DAPT wo_2h + CK1α-Deg Vs DAPT wo_2h. Volcano plot represents significant DEPs Log2 Fold Change ≥1.5, -log10 p-value ≥ 1.3. Each dot represents a biological replicate; Unpaired-T test (* P ≤ 0.05; ** P ≤ 0.01; ns = not significant); Bars represent SEM

We next tested whether it is possible to perturb N1ICD activity with a selective chemical degrader of CK1α, namely SJ3149 (CK1α-Deg) (Nishiguchi et al., 2024). Treatment with this degrader induced the decrease of the CK1α protein level (**Fig. S2A)**, and, importantly, a significant reduction of the Notch reporter signal by 25% relative to control (**Fig. 3C**). To identify the Notch signaling processes regulated by CK1α, we combined CK1α-Deg administration with NOTCH1-UltraID proximity labeling (Kubitz et al., 2022) (**Fig 3D**). We then seeded HEK293 Flp-IN cells, with mono-allelic genomic insertion of hNOTCH1-UltraID, on DLL4-coated plates to engage the receptor (**Fig. S2B**), along with the gamma-secretase inhibitor DAPT, and induce the NOTCH receptor cleavage by the DAPT washout, similarly to the NOTCH2-APEX2 approach used previously (Martin et al., 2023). We also added conditions where cells were treated with either vehicle (DMSO) or a CK1α-Deg, to test the effect of CK1α on NOTCH1 before and after ligand activation. Biotin was added to the media 15’ before collection (**Fig. 3E**). The lysates were precipitated using Streptavidin-conjugate beads and analyzed by Liquid Chromatography-Tandem Mass Spectrometry (LC-MS/MS).

The subsequent Principal Component Analysis (PCA) revealed that CK1α degradation induces a distinct shift in the global NOTCH1 C-term-proximal interactome (**Fig. 3F**). Notably, cells seeded on Fc-coated control plates (DMSO-Fc) exhibited a high baseline of N1ICD protein (**Fig. S2A**). Comparison of the DMSO-Fc control and samples inhibited by DAPT revealed a higher incidence of NOTCH1 interactions with its transcriptional co-factor MAML1, indicating substantial constitutive NOTCH1 signaling even in the absence of recombinant DLL4 ligand (**Fig. S2C, C’**). While sustained inhibition with DAPT reduced the MAML-NOTCH1 interaction relative to the 2h after DAPT washout condition (DAPT wo_2h) (**Fig. 3G, G’**), the addition of CK1α-Deg led to further decrease of NOTCH1 interactions. This was evidenced by a significant reduction in interactions with NOTCH1, NOTCH2, DTX2, MAML1, and many other proteins found in DAPT-only control condition (**Fig. 3H, H’**).

Gene Ontology (GO) analysis of the down-regulated proteins following CK1α depletion revealed a significant enrichment in transport-related signatures, specifically involving the establishment of protein localization to organelles and protein folding (**Fig. 3J, S2D**). A focused analysis of membrane and vesicle-associated proteins further identified the downregulation of COPA, Clathrin (CLTC), AP2A1 (X. Wang et al., 2008; Windler & Bilder, 2010), and multiple Rab GTPases (reviewed in Brandizzi & Barlowe, 2013; Kaksonen & Roux, 2018) (**Fig. S2F**). Together, these data suggest that CK1α depletion affects the broader NOTCH1 interactome by destabilizing the trafficking and recycling environment and the vesicle machinery required for Notch availability.

We next assessed the impact of CK1α on the dynamic assembly of the Notch active complex after DAPT washout. Heatmap visualization revealed that CK1α depletion led to a marked reduction in the interaction with NOTCH1 and MIB1 compared to the washout alone (**Fig. 3I, I’**). To probe the processes underlying the lack of interaction with Notch machinery after ligand-induced N1ICD release, we performed Gene Ontology (GO) analysis of the DAPT wo_2h + CK1α-Deg group against the DAPT wo_2h alone (**Fig. 3K, S2G**). Specifically, we observed a downregulation of key nuclear import and export machinery, including XPO1, TNPO1, TNPO3, CSE1L, and the importin KPNB1 (reviewed in Yang et al., 2023) (**Fig. S2H**). These data suggest that CK1α is also needed for the transport of N1ICD into the nucleus, required for the assembly of the N1ICD-driven transcription.

Intersection analysis across our experimental conditions helped us to further identify distinct sets of proteins exclusively expressed in a single condition or shared between specific samples (**Fig. S3**, black dots in the lower panel). We then conducted Gene Ontology (GO) analysis on these unique clusters in order to integrate the detected NOTCH1 interactome changes across the conditions. The DAPT-only NOTCH1 interactome lacked significant enrichment, while the addition of the CK1α degrader enriched proteins associated with the vesicle lumen. We identified a large set of 126 unique proteins involved in COPI vesicle interaction (H. Zhang et al., 2023), ribosome assembly, and cadherin binding 2h after DAPT washout. In contrast, treatment with CK1α-Deg shifted the protein profile toward microtubule binding, tubulin binding, and actin filament binding (**Fig. S3**). This might imply that without CK1α, the signaling machinery becomes “stuck” in proximity to the plasma membrane, tethered to the cytoskeletal framework rather than transitioning into the active transport phase. Furthermore, intersection analysis showed that DAPT and wo_2h share a core signature in carrier activity and transport, which is completely lost in their CK1α-Deg counterparts. These findings indicate that CK1α is heavily involved in the transport of the N1ICD pool towards the nucleus, and possibly in its ability to interact with other proteins as such. Given the observed loss of interactions in CK1α absence affects both the NOTCH1 pre- and post-activation state, it distinguishes itself from the degradation-related effects of GSK3β or CDK1/2 as well as the trafficking-sorting activity of PKCζ-mediated NOTCH1 phosphorylation (Sjöqvist et al., 2014).

### CK1 phosphorylates several S/T of N1ICD in the N-terminal ankyrin repeat region

To examine whether the effects of CK1α on NOTCH1 are direct or not, we tested possible interaction of the two proteins. Co-transfection of HEK293T cells with HA-tagged NOTCH1 and 3xFlag-tagged CK1α resulted in successful co-immunoprecipitation of the CK1α-NOTCH1 complex (**Fig. 4A**), which, together with dominant-negative effect of the kinase-dead CK1α (**Fig. 2E**), suggested that NOTCH1 can be a direct phosphorylation target of CK1α. We thus took advantage of kinase domain sequence similarity between CK1α and CK1e (Knippschild et al., 2005) and performed *in vitro* phosphorylation assay followed with LC-MS/MS of N1ICD-ANK core (N1ICD aa 1872-2126) (Choi et al., 2012) with CK1 core (CK1e aa 1-301) (Harnoš et al., 2019). Mass spectrometry analysis detected phosphorylation at seven S/T sites, specifically S1889, S1891, T1897, S1900, T1930, S1951, and S1970 of human N1ICD (**Fig. 4B, C**). Positions S1889, S1891, T1897, S1900 in ANK repeat 1 were the most prevalent, while T1930 and S1951, flanking ANK2, and S1970 in ANK3 were present with lower frequency (**Fig. S4A**). Of note, none of the sites except for T1897 and S1900, phosphorylated by CK2 (Ranganathan et al., 2011) were previously shown to be phosphorylated (Antfolk et al., 2019).

**Figure 4:**
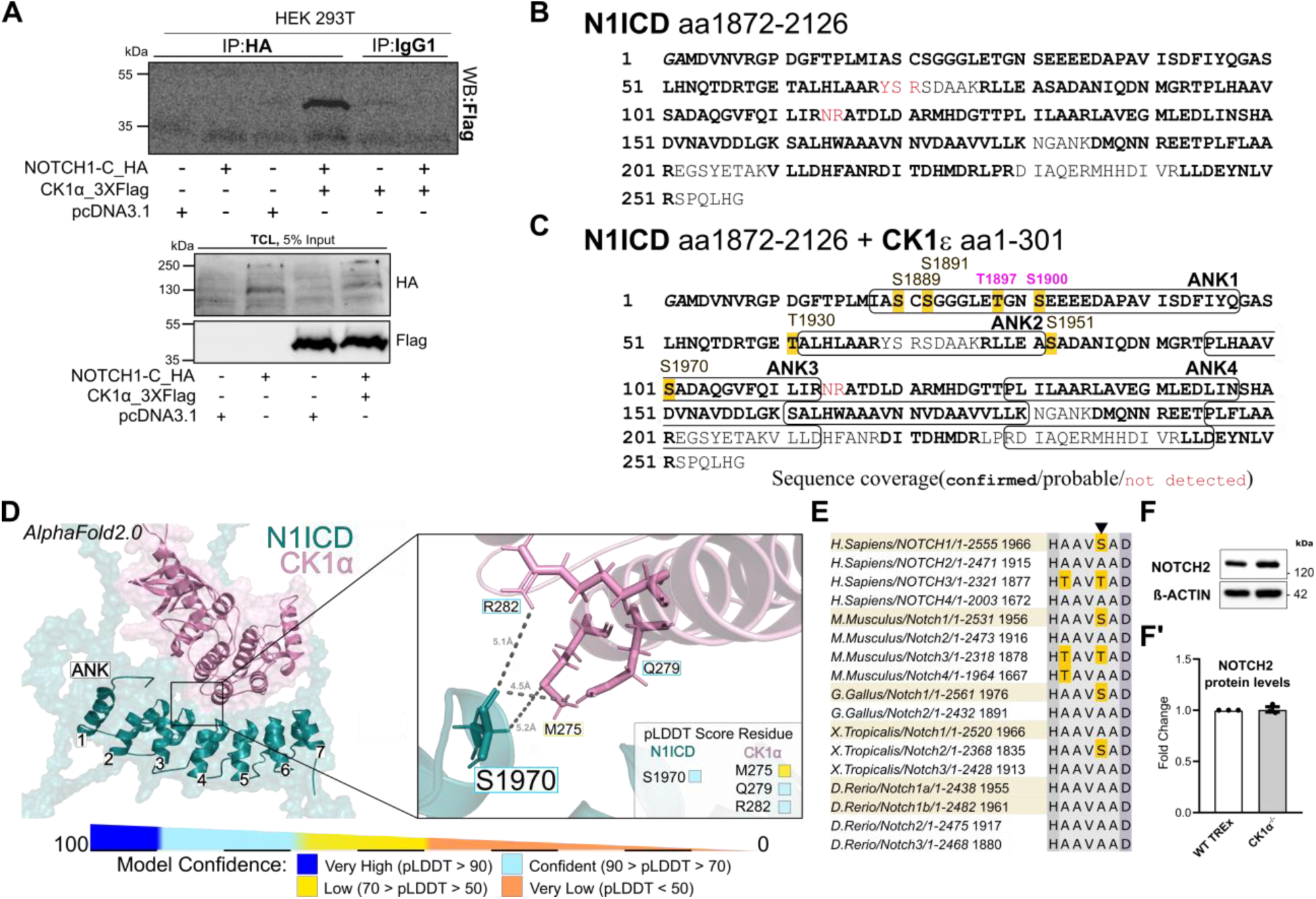
CK1α facilitates the proper assembly of the Notch transcriptional machinery via Serine 1970. (**A**) Co-immunoprecipitation with HA antibody or IgG1 control, followed by western blot of Flag in WT HEK 293T transfected with NOTCH1-HA and 3xFlag-CK1α. (**B-C**) in vitro phosphorylation assay followed by LC-MS/MS of (B) N1ICD-ANK core (aa1872-2126), and (C) N1ICD-ANK core + CK1ε (1-301). (**D**) N1ICD-CK1α AlphaFold2.0 modelling prediction. (**E**) Multiple amino acid sequence alignment of NOTCH receptors. (**F**) Western blot of NOTCH2, in WT TREx and CK1α^-/-^ cells with (**F’**) protein quantification. β-ACTIN was used for normalization. (Annotation of the ANK repeats in C is based on Ehebauer et al., 2005. pLDDT, predicted local distance difference test.)

We next employed the *in silico* AlphaFold2.0 prediction (Jumper et al., 2021) to further test if full length CK1α can interact with any of the newly identified phopsho-sites. First, we confirmed the reliability of the approach using the existing crystal structure of N1ICD-RBPJk-MAML (Choi et al., 2012), yielding a cumulative confidence score of 81.1, with a balanced confidence level distribution between residues of MAML and N1ICD (**Fig. S4B, C**). Furthermore, as a negative control, we tested N1ICD interaction with Notch-unrelated mitochondrial protein MPPB, resulting in a cumulative score of 51.6 and uneven distribution of confidence between residues of N1ICD (23.15) and MPPB (85.83) (**Fig. S4D**).

After this validation, we used AlphaFold2.0 for the prediction of the N1ICD-CK1α interaction (**Fig. 4D**). Our analysis revealed a candidate site for the direct interaction of N1ICD-CK1α with a prediction score of 74.9 (balanced), and a low confidence prediction for the N1ICD-AXIN1 46.7 (uneven) (**Fig. S4F**). Of note, N1ICD-CK2 prediction (**Fig. S4G**) fit the phosphorylation motif reported by Ranganathan and colleagues (Ranganathan et al., 2011). Interestingly, the S1970 residue is Alanine in non-avian homologs and became Serine in avian/mammal NOTCH1, not present in other NOTCH paralogs (**Fig. 4E**). We thus hypothesized that if the S1970 is NOTCH1-specific CK1α target, NOTCH2 should be unaffected in *CK1α*^-/-^ cells. Indeed, we found the amount of NOTCH2 protein in *CK1α*^-/-^ cells to be at the same level as in WT TREx cells (**Fig. 4F, F’**). Based on these results, we recognized the S1970 residue in N1ICD to be a novel NOTCH1-specific CK1α target (**Fig. 4B-D**), which we investigated further.

### NOTCH1 S1970 is critical for N1ICD transcriptional activity *in vivo*

Phosphorylation of NICD has been previously shown to be essential for NICD-CSL mediated transcription (Foltz & Nye, 2001), but details of the mechanism are unknown. At the same time, CK2-mediated phosphorylation of S1901 decreases MAML-NICD affinity and reduces the NICD transcriptional output (Ranganathan et al., 2011). To investigate whether CK1α regulates NOTCH1 activity through phosphorylation at S1970, we generated non-phosphorylatable (S1970A) and phosphorylation-mimicking (S1970D) variants of NOTCH1 and corresponding N1ΔE constructs (S1960A/D in mouse) (**Fig. 5A**). The mN1ΔE-S1960D variant resulted in comparable level of Notch reporter activation as the WT N1ΔE in *CK1α*^-/-^ cells, while the mN1ΔE-S1960A achieved less than half of the WT mN1ΔE activity (**Fig. 5B**), indicating that some level of mN1ICD-S1960A activity is maintained when overexpressed. To further assess if the S1970 is the substrate for the CK1α kinase activity, we employed again the Notch reporter assay in N1-4 KO cells seeded on a DLL4-coated plate. We observed that CK1α-KD overexpression significantly reduced CK1α-NOTCH1-dependent transcriptional activity, while the phospho-mimic NOTCH1 S1970D as well as the S1970A mutants remained unresponsive to CK1α-KD (**Fig. 5C**).

**Figure 5:**
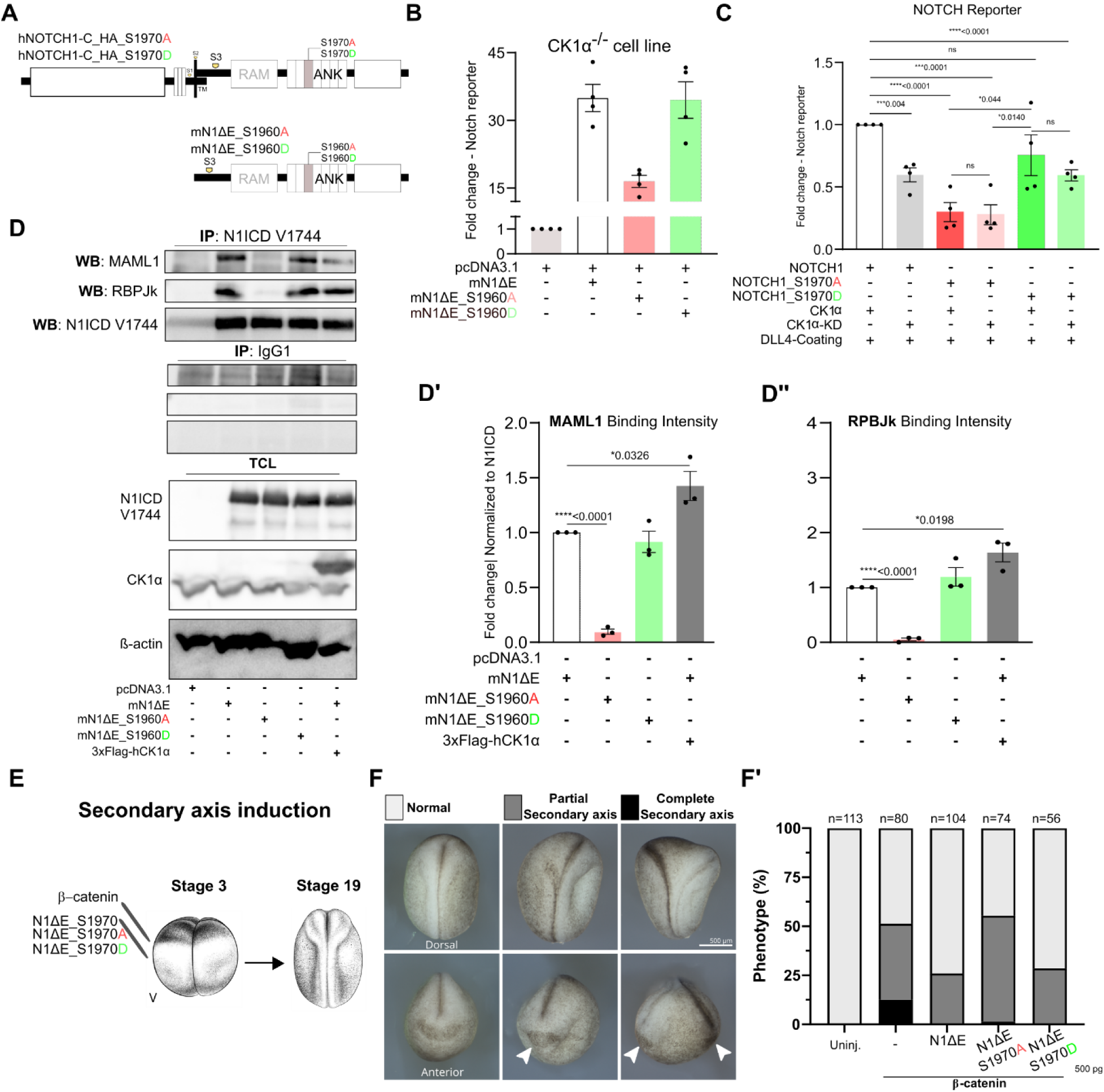
CK1α facilitates the proper assembly of the Notch transcriptional machinery via Serine 1970. (**A**) Scheme of NOTCH1-S1970A/D and mNIΔE-S1960A/D generated plasmids. (**B**) Luciferase Notch reporter assay in CK1α^-/-^ cell transfected with pcDNA3.1 and WT, S1960A/D mNIΔE plasmid. (**C**) Luciferase Notch reporter assay in N1-4KO cells on Fc-DLL4 coated plates, transfected with NOTCH1, S1970A/D, and either CK1α or CK1α-KD plasmid. (**D**) Co-immunoprecipitation with N1ICD Val1744 antibody or IgG1 control, followed by western blot of MAML1, RPBJk, and N1ICD, in N1-4KO cells transfected with WT, S1960A/D mNIΔE, and 3xFlag-CK1α, with quantification of (**D’**) MAML1 and (**D’’**) RBPJk intensity. Each dot represents a biological replicate; Unpaired-T test (* P ≤ 0.05; ** P ≤ 0.01; *** P ≤ 0.001; **** P > 0.0001; ns = not significant); Bars represent SEM. (**E**) Schematic representation of the secondary axis assay. hβ-catenin alone or together with mNIΔE WT, S1960A/D mRNA was injected into both ventral blastomeres of 4-cell stage Xenopus laevis embryos. (**F**) Representative embryos showing normal morphology, partial secondary axis, and complete secondary axis phenotypes. Complete secondary axes are indicated by white arrowheads. Dorsal and anterior views are shown. Scale bar, 500 µm. (**F’**) Quantification of secondary axis phenotypes. Total number of analyzed embryos per condition is indicated above each bar (n = 56–113, 3 independent biological replicates).

To assess the impact of NOTCH1 S1970 variants on interaction with the core Notch transcriptional partners MAML, and RBPJk (reviewed in Bray & Gomez-Lamarca, 2018; Nam et al., 2006), we expressed either wild-type or mutant forms of N1ΔE in cells and performed co-immunoprecipitation using N1ICD-specific antibody (Val1744). The S1960A substitution markedly reduced the interaction of N1ΔE with both the endogenous MAML1 and RBPJk, whereas the phospho-mimic S1960D maintained binding comparable to the wild-type (**Fig. 5D-D’’**). Overexpression of CK1α further enhanced N1ICD association with both co-factors, consistent with a phosphorylation-dependent stabilization of the transcriptional complex (**Fig. 5D-D’’**).

We next used a well-established readout of Wnt-dependent secondary axis induction in *Xenopus laevis* embryos (McCrea et al., 1993; Sokol et al., 1991). This ectopic axis induction has been previously shown to be impaired by NICD through interference with β-catenin activity (Acosta et al., 2011), and was used here to functionally test the requirement of S1970 phosphorylation *in vivo*. First, we injected hβ-CATENIN into equatorial region of both ventral blastomeres of a 4-cell stage embryo (**Fig. 5E**). In β-catenin–injected embryos, a partial or even complete secondary axis was observed (**Fig. 5F, white arrowheads**). When we co-injected WT mN1ΔE or the mN1ΔE-S1960D, we observed a rescue - prevention of the secondary axis development, while the mN1ΔE-S1960A injection failed to do so (**Fig. 5F, F’**). Together, these results identified NOTCH1-specific S1970 integrity as a requirement for N1ICD transcriptional complex assembly *in vivo*.

### Phosphorylation of NOTCH1 S1970 does not affect N1ICD nuclear localization, but alters N1ICD ANK2-3 conformation via intrinsic interactions with NOTCH1 R1937 and R1962

Our data indicate the NOTCH1/N1ICD “interactivity” both in cytosol and in nucleus depends on the presence or absence of CK1α. We next aimed to test if the S1970 is also important for the NOTCH1 subcellular distribution and the interaction with CK1α itself.

We hypothesized that disrupted assembly of the MAML1-RBPJk-NOTCH1 S1970A (**Fig. 5D**) may be caused by the N1ICD failure to reach the nuclei, especially given the lack of interactions with nuclear transport proteins (**Fig. 3K**). We thus examined the subcellular distribution of NOTCH1 and CK1α by confocal microscopy in N1-4KO cells. Cells were seeded on DLL4-coated coverslips, treated with DAPT to prevent NOTCH1 cleavage and accumulation at the plasma membrane or following a 2-hour DAPT washout to allow receptor release and N1ICD nuclear translocation. Strikingly, our quantitative colocalization analysis revealed that the colocalization score of NOTCH1-S1970A with nucleus (Hoechst) was significantly higher than NOTCH1-S1970D following the DAPT washout (**Fig. S5B-C**).

The high degree of colocalization of NOTCH1 and CK1α in control condition (Fisher’s z’ transformation 0.72) (**Fig. S5A, 6A’**) showed their proximity indicated by co-immunoprecipitation (**Fig. 4A**). The colocalization score of WT NOTCH1 and CK1α had an increasing trend after DAPT treatment (Fisher’s z’ transformation from 0.72 to 0.89), whereas the CK1α - NOTCH1-S1970A/D score was significantly lower (Z-scores of 0.41 and 0.67) (**Fig. 6A, A’**). However, this difference was lost again after the DAPT washout (**Fig. S5B, 6A’**), indicating that the pre-S3 cleavage events may be a hotspot for the CK1α-NOTCH1 S1970 interaction. While possibly masked to some extent by the overexpression system, our microscopy data thus do not indicate any localization defects that could prevent the ternary complex formation with MAML1 and RBPJk.

**Figure 6:**
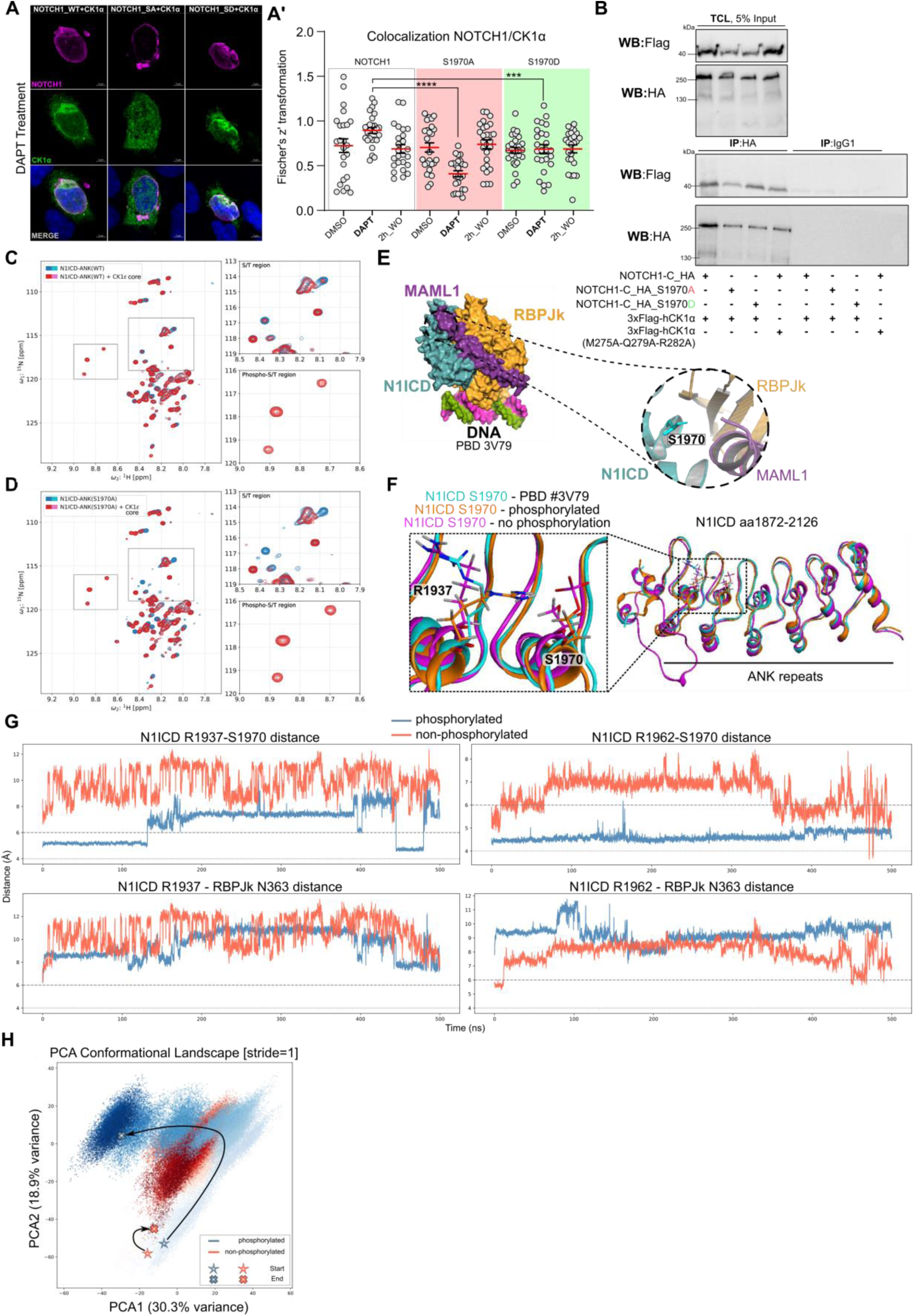
S1970 N1ICD phosphorylation does not disrupt nuclear localization but regulates N1ICD ANK repeat conformation Via NOTCH1 R1937 interaction. (**A**) Representative confocal images (MIP) of DAPT-treated cells transfected with NOTCH1-S1970A/D and 3xFlag-CK1α. (**A’**) Fischer’s z’ quantification of NOTCH1 (WT, S1970A, S1970D) and CK1α colocalization. N1-4KO cells seeded on a Fc-DLL4 coated plates were treated with DAPT. Scale bars: 5µm. Ordinary One-way ANOVA; **\*\*\***P < 0.001, **\*\*\*\***P < 0.0001. (**B**) Co-immunoprecipitation with HA antibody or IgG1 control, followed by western blot of Flag and HA in WT HEK 293T transfected with WT, S1970A/D NOTCH1, and 3xFlag-CK1α or 3xFlag-CK1α M275A-Q279A-R282A. (**C,D**) 1H-15N HSQC spectra visualization of NMR measurements following in vitro phosphorylation of (C) WT-N1ICD-ANK or (D) S1970A-N1ICD-ANK with CK1ε core. (**E)** Crystal structure (PDB 3V79), showing relative location of N1ICD S1970 to MAML1 and RPBJk. **(F,G**) Molecular dynamics simulations of the N1ICD ANK repeats. (F) Structural overlays of unphosphorylated (magenta) and pS1970 (orange) states against PDB 3V79 (cyan). (G) Plot of the distance between Arg1937, Arg1962 and Ser1970 NOTCH1 residues and RBPJk N363. **(H**) PCA of un-and phosphorylated complex dynamics showing a phosphorylation-dependent shift. The star and cross represent the 500ns simulation (start/end). Scale bars: 5µm. Ordinary One-way ANOVA; **\*\*\***P < 0.001, **\*\*\*\***P < 0.0001.

To explore further the nature of CK1α and N1ICD interaction that could mechanistically uncover the role of S1970 phosphorylation, we co-transfected the HEK293T cells with 3xFlag-tagged CK1α and either WT or NOTCH1 S1970D/A variants and performed a co-precipitation using αHA-antibody. Both NOTCH1 S1970D/A precipitated with CK1α (**Fig. 6B**), which was consistent with the microscopy results. We next tested the binding properties of WT NOTCH1 and CK1α motif M275-Q279-R282, which was predicted to interact with NOTCH1 S1970 (**Fig. 4D)**. As CK1α M275A-Q279A-R282A mutant also precipitated together with NOTCH1 (**Fig. 6C**), it suggests that the NOTCH1-CK1α interaction occurs through other residues.

If that were the case, S1970 would likely not act as a “seed” residue required for initiation of the phosphorylation of the preceding sites. To test this hierarchical dependency in solution, we monitored the *in vitro* phosphorylation of the N1ICD aa1872-2126 by CK1ε core using real-time 1H-15N HSQC NMR. Very few peaks are present in the NMR spectrum, presumably corresponding to the flexible, N-terminal segment with ANK1 and not the rigid core. Upon the addition of the CK1ε core (i.e., kinase domain), we tracked the phosphorylation kinetics over 14 hours, observing the progressive appearance of three (possibly four) distinct phospho-S/T peaks (**Fig. 6C**). The observed phosphorylations should correspond to the S/T residues reported by MS residing in the disordered region N-terminally to ANK repeats. To determine if S1970 drives the observed phosphorylation events, we performed comparative NMR kinetics using an S1970A variant. This was not the case, as the WT N1ICD-ANK and S1970A N1ICD-ANK spectra were nearly identical (**Fig. 6D**).

Examination of the crystal structure of the N1ICD-RBPJk-MAML1 ternary complex (PDB 3V79, formed with non-phosphorylated N1ICD from *E. coli*) (Choi et al., 2012) revealed that S1970 lies at the periphery of the MAML1 and RBPJk binding surfaces (**Fig. 6E**). This suggests that the S1970 might affect the complex conformation, rather than its assembly, similarly to the reported CDK5-mediated phosphorylation impact on interactions within the ANK repeats of TRPA1 (Hall et al., 2018). We thus performed molecular dynamics simulations of the N1ICD-RBPJk-MAML1 ternary complex before and after S1970 phosphorylation over 500 ns. Strikingly, upon phosphorylation, the S1970 phosphate group becomes stably coordinated by an intramolecular salt bridge between ANK2-3 involving R1937 and R1962. This electrostatic “clamping” sequesters these arginines and disrupts a key inter-subunit contact, breaking the persistent hydrogen bond between NOTCH1 S1970 and RBPJk N363 observed in the unphosphorylated state (**Fig. 6F, G**).

Interestingly, the dynamic cross-correlation and principal component analyses indicate that S1970 phosphorylation weakens the coupling between N1ICD and RBPJk, shifting their relationship from co-correlated to anti-correlated motion **(Fig. 6H, S5D**). Concurrently, the RBPJk and MAML subunits become more tightly coupled, moving as a semi-rigid unit decoupled from NOTCH. Ultimately, this phosphorylation event induces major conformational reorientations, captured largely along the principal axis of motion, driving significant structural shifts in the RBPJk interface region and the MAML C-terminus. While not offering a direct explanation, this complex rearrangement could affect the conformation of the flexible, RAM domain containing, N1ICD N-terminus, which is crucial for full transcriptional activity as recently shown by Cook and colleagues (Cook et al., 2025). Together, these results suggest that CK1α primarily associates with NOTCH1 in the cytoplasm, mediating phosphorylation at S1970 which acts as intrinsic N1ICD modulator of subsequent receptor interactions in the cytoplasm and transcriptional activity in the nuclei.

## DISCUSSION

Precise regulation of Notch receptor signaling is fundamental for cell differentiation and maintenance, yet the proposed interaction between Notch and the components Wnt/β-catenin pathway remain poorly characterized (Hayward et al., 2008; Turetti et al., 2026). In this study, we describe how lack of individual Wnt/β-catenin components affects the steady state levels of the active, S3 cleaved, NOTCH1 intracellular domain. Our data confirmed previously described positive effect of GSK3β (Foltz et al., 2002b; Guha et al., 2011), and the side-by side comparison brought new evidence of baseline modulation of N1ICD levels by AXIN1 and to a lesser extent also E3 ligases RNF43 and ZNRF3. However, the key discovery of this work is the previously unrecognized CK1-mediated phosphorylation of multiple S/T sites in the NOTCH1 intracellular domain ANK domain. While CK1α is a canonical negative regulator of the Wnt signaling via the destruction complex, our study introduces it as a novel positive regulator of the NOTCH1 activity regardless of the β-CATENIN level or WNT3a activation, thus likely acting outside of the “canonical” Wntch concept. Crucially, we demonstrate that CK1 kinase activity-dependent synergy with N1ICD potentiates Notch target gene expression *in vivo.* We further identify several new phosphorylation sites in the N1ICD ANK region, bringing evidence of S1970 phosphorylation as key, functional NOTCH1 ANK3-specific modification that, likely via intrinsic changes in conformation, promotes N1ICD-RBPJk-MAML1 transcriptional complex activity, as we demonstrate using the axis duplication-rescue assay in *Xenopus*.

Several recent studies have recently assessed the proximal proteome of multiple Wnt/β-catenin components and Notch receptors (reviewed in Turetti et al., 2026). Our proteomics experiments expand the data further by incorporating the PROTAC-mediated degradation of CK1α. Interestingly, the key observed effect on the NOTCH1 C-terminus was a widespread loss of interactivity or mobility. Among the significantly downregulated proteins after CK1α depletion stands out the Annexin family (ANXA1/2/5/6). Annexins are established regulators of membrane-associated actin dynamics and endocytic pathways and have recently been identified as core components of the human Notch1 interactome (Bian et al., 2023; G. Shao et al., 2024; Singh et al., 2026). The loss of annexin-mediated scaffolding could explain the disrupted sequestration of NOTCH1 within the trafficking pipeline. The trafficking defect is further underscored by the fact that TMEM214 (C. Li et al., 2013), TMEM131 (Z. Zhang et al., 2020), and the clathrin-regulator GAK (Kanaoka et al., 1997) were among the top 10 most downregulated proteins in the DAPT+CK1α-Deg comparison condition. These findings highlight a collapse of the machinery required for NOTCH1 membrane proximity and intracellular transport. Consistent with a direct regulatory role, CK1α as well as ANXA2 and CAV1 proteins were identified within the NOTCH1 precipitate under physiological flow in the polarized membrane microdomains of primary human dermal blood microvascular endothelial cells (Singh et al., 2026), reinforcing the default role of CK1α in the NOTCH1 life cycle.

Our proximity assay revealed close relationship between NOTCH1 and other Notch receptors (NOTCH2/3). These findings are consistent with the recently suggested requirement for NICD homo- and hetero-dimerization in driving transcriptional activation (Gazdik et al., 2024) and could explain the seeming discrepancy of the proximity assay, concluding a disrupted nuclear transport of N1ICD in the absence of CK1α, and confocal microscopy results, indicating the N1ICD S1970A presence in the nuclei. The key difference is in usage of WT cells with presence of NOTCH 2 and 3 along the NOTCH1 that compete for the binding of the components that showed lower affinity to the NOTCH1 without phosphorylation the CK1α-Deg condition; on the other hand, this cannot happen when we tested the NOTCH1 S1970A/D variants in the NOTCH1-4KO cells, allowing for transport to the nuclei, likely less dynamically. While speculative at this point, such hypothesis can be experimentally tested in the future.

Our data also point towards new self-regulatory feedback between N1ICD, CK1α, and MIB1. After Notch activation, we observed a marked reduction in MIB1-NOTCH1 proximity upon CK1α depletion suggesting the Notch activation and CK1α stability are interlinked. Interestingly, MIB1 has recently been identified as an E3 ubiquitin ligase that targets CK1α for degradation (X. Li et al., 2025). Our findings thus point toward a self-limiting mechanism: active NOTCH1 may recruit MIB1 to degrade CK1α, reducing the available N1ICD pools and ensuring a transient signaling pulse.

Through the *in vitro* phosphorylation assay using the CK1ε core followed by LC-MS/MS, we identified five novel and two previously reported S/T phospho-sites within the N1ICD (Antfolk et al., 2019; Ranganathan et al., 2011). Using AlphaFold modelling we corroborated these findings, suggesting that N1ICD S1970 is targeted specifically by CK1α, which we confirmed to be essential for ternary complex assembly and transcriptional activity. Notably, previous LC-MS/MS studies of HEK293T cells transfected with hN1ICD-GFP illustrate that detecting the phosphorylation in the ANK region is problematic, as it recognized the S1900 (Carrieri et al., 2019), but failed to detect the other CK1/CK2-targeted T1897 site (Ranganathan et al., 2011), or other phospho-sites reported in this study, including the S1970. While the direct phosphorylation of S1970 by full-length CK1α remains to be confirmed, our results highlight S1970 as one of the functional switches through which NOTCH1 activity is mediated.

We used molecular dynamics simulations to gain more insight into the conformation of N1ICD-ANK-RBPJk-MAML1 after N1ICD S1970 phosphorylation. Our modelling indicates that the chemical state of S1970 governs an intramolecular salt bridge between R1937 and R1962, within the ANK repeats. Our data indicates that following the phosphorylation, the trimeric complex of N1ICD-ANK-RBPJk-MAML1 rearranges quite dramatically, possibly to facilitate a better interaction of the intrinsically disordered N-terminus containing the RAM domain, which was recently proposed to wrap around the MAML1-RBPJk complex (Cook et al., 2025). The fact that N1ICD S1970A variant, unlike the non-phosphorylated N1ICD-ANK used for the trimeric complex crystal structure (Choi et al., 2012), fails to stably recruit the core co-factors MAML1 and RBPJk, indicates that the N-terminal part of the N1ICD might be indeed affected by the PTMs, such as the S1970 phosphorylation. This aligns with other PTMs in the same region, such as Src-mediated phosphorylation, which inhibits MAML1 binding (LaFoya et al., 2018), and hydroxylation at N1945, which regulates signaling dynamics (Ferrante et al., 2022). Notably, the resulting drop in NOTCH1 is reflected not only at the protein levels but also in a significant reduction of *NOTCH1* transcript, consistent with reports that suppressing Notch transcriptional activity triggers a negative feedback loop on *NOTCH1* expression (Chen et al., 2017), suggesting that the CK1α-S1970 axis is required for maintaining the signaling pool at both the transcriptional and post-translational levels.

Identification of S1970 is particularly interesting given its absence in NOTCH2, which we found to be unaffected by CK1α loss. NOTCH1 specificity extends to other identified phosphoresidues; S1891, T1897, S1900, and S1951 are all unique to NOTCH1, potentially facilitating isoform-specific regulation. S1889 is conserved across NOTCH1/2/3, while T1930 is present in NOTCH1/3, and 4. Of note, our data to some extent challenge the interpretation of the genetic from N1-N2ICD swapping experiment in mice (Z. Liu et al., 2015), however it should be noted that CK1α is widespread and thus likely capable of phosphorylation of the N1ICD also in the usually Notch2-expressing tissue. We demonstrate biological relevance of this S1970 by our *Xenopus* experiments, where the mN1ΔE S1960A failed to rescue the secondary axis formation induced by β-catenin overexpression. It has been previously established that the NICD suppresses β-catenin effect by destabilizing maternal β-catenin (Acosta et al., 2011). Our data reveal that this interaction depends on the presence of the N1ICD S1970 residue. Notably, despite using a 97.5% lower concentration of the N1CD mRNA than the levels reported by Acosta and colleagues, we observed that both WT NICD and the phosphoylation-mimicking variant S1970D successfully inhibited axis duplication. In contrast, the N1ICD S1970A mutant was largely unable to suppress the secondary axis, failing to rescue the normal embryonic patterning. This indicates that even at very low doses, the integrity of the N1ICD S1970 residue is a prerequisite for the functional interference between Notch and β-catenin signaling *in vivo*, possibly due to the loss of N1ICD interactivity, stemming from the altered conformation of the ANK2-3 repeats.

Of note, our results demonstrate that while AXIN1/2 loss leads to robust hyperactivation of Wnt signaling, it has only a partial effect on NOTCH1/NICD protein levels. The observation that AXIN1 overexpression decreases the Notch signaling response aligns with the hypothesis that Notch may operate in a parallel pathway that cooperates with AXIN1 independently of the canonical destruction complex (Hayward et al., 2006). This suggests that while both proteins are core members of the Wnt destruction complex, they possess different roles in the “Wntch” signaling.

Our study has several limitations that should be mentioned; proximity-based approaches have inherent caveats that must be considered. First, the approach is sensitive to overexpression artifacts, and the absence of certain proteins in our dataset, such as RBPJk and CK1α, does not necessarily imply their absence within the physiological complex, but rather that they might be below the level of detection or outside the effective labeling radius in our specific experimental setup due to C-terminal position of the UltraID enzyme. Second, our interactome was established in Flp-in HEK293T cells. An inevitable limitation is that cell-type-specific partners may not be identified. Establishing how these molecular signatures shift in other relevant models will be a critical step toward a comprehensive understanding of NOTCH1 signaling dynamics. On the other hand, we observed both synergistic effects of CK1α and N1ICD on *Xenopus* Notch target *Hairy2* as well as S1970 dependent rescue of the axis duplication assay.

A significant consideration emerging from our study is the potentially redundant or synergistic roles of CK1α with the Casein Kinase 1 family paralogs (Cheong & Virshup, 2011). Recent Proximity-Dependent Biotinylation (mini-Turbo) datasets revealed the presence of NOTCH1-2-3 in the interactomes of CK1γ isoforms (1-2-3) (Agajanian et al., 2022). This is complemented by findings that CK1δ/ε also associates with full-length NOTCH1 (Bian et al., 2023). CK1ε overexpression in HCT116 and HT29 has been shown to increase transcript levels of *HES1* and *HES2 (Z. Wang et al., 2021)* in an Amino-terminal enhancer of split (AES)-dependent manner, which was shown to create nuclear condensates with NICD (Gazdik et al., 2025). Pharmacological inhibition of CK1ε has been shown to effectively reduce Aβ formation in N2a cells without inhibiting Notch cleavage, even though this experiment did not assess downstream signaling activity (Flajolet et al., 2007). These observations align with our findings, suggesting that the CK1 family may potentiate NOTCH1 signaling competence at the level of the intracellular domain rather than through the modulation of receptor degradation.

Finally, CK1γ3 overexpression significantly increases Notch reporter activity, suggesting that the “licensing” of Notch signaling might be distributed function across the CK1 family. This is supported by evolutionary data from Drosophila, where the CK1γ homolog Gilgamesh (Gish) regulates the Notch pathway independently of canonical β-catenin signaling (S. Li et al., 2020; Swarup et al., 2015). Together, our findings add a new layer to the NOTCH1-specific life cycle, identifying a default CK1-maintained PTM and a previously unrecognized residue that is essential for the receptor to reach its full signaling potential. Acting irrespective to WNT3a activation, CK1α acts most likely outside of the joint Wntch concept, inviting for further, focused probing of this signaling concept. By bridging the gap between trafficking scaffolding and transcriptional licensing, the CK1α-S1970 axis opens new aspect of Notch receptor biology; that ought to be considered when developing novel CK1 or NOTCH1-specific therapeutics.

## MATERIALS AND METHODS

### Cell Culture

HEK293T, N1-4KO cell (kindly gifted by Andersson’s lab) and HEK T-REx™-293 cells (all the Wnt Knockout cell lines were kindly gifted from Vítězslav Bryja’s lab) were cultured in DMEM + GlutaMAX™ (Gibco, #31966047) supplemented with 10% FBS (Thermo Scientific, #A5256701), 1% penicillin – streptomycin (Gibco, #15140122). Cells were maintained at 37°C, in 5% CO2 in a humidified HERACell VIOS 250i incubator and were routinely tested for Mycoplasma contamination.

### Generation of HEK Flp-In™ hNOTCH1-UltraID expressing cells

To generate the hNOTCH1-UltraID stable expressing cell lines, HEK Flp-In™ cells were utilized according to the manufacturer’s instructions. Briefly, the ultraID sequence was amplified from the pSF3-ultraID template (Addgene, Plasmid #172878) using Q5 High-Fidelity DNA Polymerase (New England Biolabs, #M0491S). Primers were designed to incorporate an HA-tag and an XhoI restriction site at the 5’ terminus, and BclI and XhoI sites at the 3’ terminus (FT_F11+FT_R19). The resulting PCR product was digested with XhoI and ligated into the pcDNA3.1-hNOTCH1 plasmid. Correct insertion was verified via PCR using specific flanking primers (FT_F13+FT_R18) and Sanger sequencing. Subsequently, the hNOTCH1-HA-ultraID fragment was subcloned into the pcDNA5/FRT vector using EcoRI and BclI restriction enzymes. To ensure efficient BclI digestion, plasmids were propagated in a Dam-deficient (Dam^-/-^) bacterial strain to prevent site-specific methylation. For genomic integration, pcDNA5/FRT-hNOTCH1-HA-ultraID was co-transfected with the pOG44 Flp-recombinase expression plasmid at a 1:9 ratio into HEK Flp-In™ cells. Stable clones were isolated via single-colony selection and validated for both correct protein expression and biotinylation functionality.

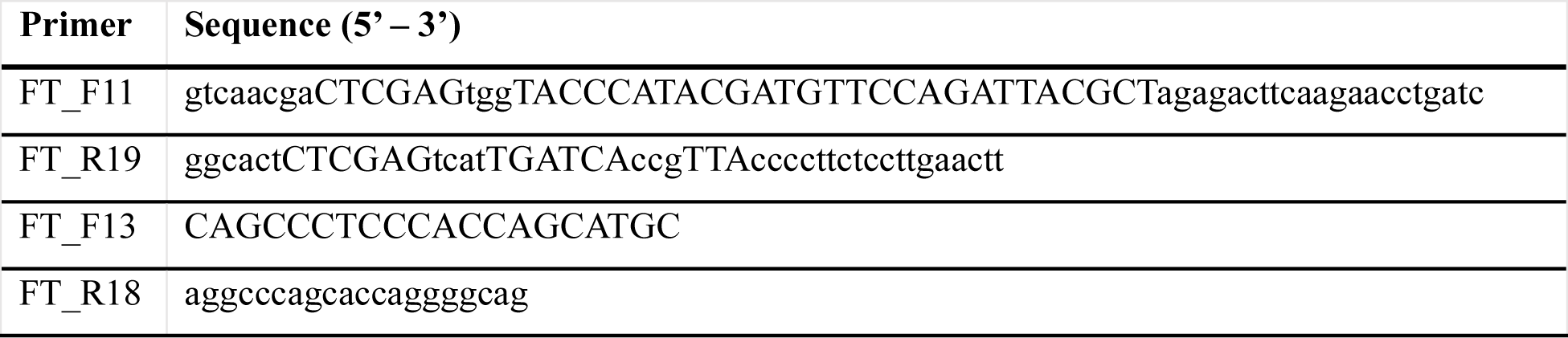

### Immunostaining

Coverslips were washed in 100ml of 1M HCl for 16h at 60C. Then washed two times with dH2O and two times with ddH2O. Finally washed in 96% Ethanol and dried between sheets of Whatman filter paper and sterilized by UV for 20 minutes. Acid washed coverslips were coated in 0,1% gelatine (in 1X PBS) before cells were seeded for immunofluorescence. Solution of gelatine was heated to 37°C. Then 200µl was applied onto each coverslip in a 24 well plate and incubated at RT in a sterile environment for about 15 minutes. After incubation time, full volume was aspirated and left to semi-dry for about 10 minutes. Cells were kept in an incubator for 16 hours after seeding, transfected, and after 4 hours medium was changed. If cells were treated with DAPT/DMSO (20µM) it was added with a new substrate and after 24 hours treatment was applied again for an additional 2 hours. 24 hours after transfection cells were washed with 1xPBS. Samples were fixed with freshly made 4% paraformaldehyde in 1xPBS. Cells were washed 3 times with 200 μl of PBST, blocked with 200μl of 10% BSA PBT(A), and incubated with primary antibodies overnight at 4°C in PBTA with 3% BSA. Samples were washed 3x with PBST for 10 minutes, then incubated at room temperature in secondary antibodies diluted 1:500 in PBT(A) 3% BSA. Samples were washed 2x with 200 μl of PBST. Third wash was done with Hoechst 33342, diluted 1:100 in PBST to the total volume of 200 μl and incubated for 10 minutes at room temperature. Then samples were washed 3x with PBS. Glass slides were cleaned with 70% EtOH and a drop of Fluoroshield mounting media applied.

### Microscopy Images Acquisition

Images were acquired using fluorescence inverted confocal microscope Zeiss LSM 880 with ZEN Black software. Maximum Intensity Projections (MIP) were created in ZEN Blue.

Plan-Apochromat, Oil, 63x objective was used for acquiring the images. All images were captured as Z-stacks, thickness 0.32 µm per focal plane. Using a 2048×2048 image size, 16-bit depth. Scan direction – Unidirectional, averaging 2, 0.52 us pixel dwell time, 15.19s Frame time (for 3 channels), Scan Zoom 1.0. Pinhole 0.91A-0.92 AU – Airy was set to 1. Hoechst 33342 channel was captured using 405 nm Laser 30mW. Laser power was set to 4%, Detector Gain from 550-630. Detector digital gain to 1.00. Detection Wavelength 410-499. MBS: Plate, MBS-InVis: -405., MainBeamSplitterNonDescanned: Rear. AF488 channel was captured using Argon Laser 488 nm 25 mW. Laser power was set to 3%, Detector Gain from 470-570. Detector digital gain to 1.00. Detection Wavelength 499-562. MBS:488. MBS-InVis: Plate, MainBeamSplitterNonDescanned: Rear. AF594 staining channel was captured using 561 nm Laser 20 mW. Laser power was set to 6%, Detector Gain from 500-600. Detector digital gain to 1.00. Detection Wavelength 579-633. MBS:488/561/633. MBS-InVis: Plate, MainBeamSplitterNonDescanned: Rear. AF647 staining channel was captured using 633 nm Laser 5 mW. Laser power was set to 4%, Detector Gain from 420-520. Detector digital gain to 1.00. Detection Wavelength 638-755. MBS:488/561/633. MBS-InVis: Plate, MainBeamSplitterNonDescanned: Rear.

Data Analysis

Microscopy data were analysed in FIJI using JaCOP Biop extension (Bolte & Cordelières, 2006) – created by Bioimagining and Optics Platform (PTBIP) group from EPFL University. Available at https://wiki-biop.epfl.ch/en/ipa/fiji/update-site. ROI for each cell was selected, Automatical Otsu Thresholding was applied for each channel and colocalization result was obtained by Pearson’s correlation coefficient (r). Values for Pearson’s r were transformed to Fisher’s Z-score by using this formula: z’ = .5[ln(1+r) – ln(1-r)] to allow for statistical analysis. Statistics was performed in GraphPad Prism 11. Student’s t-test for comparison between two groups or one-way ANOVA (with reference to the first sample in the given group) for multiple comparisons was applied. P-values: >0.05 = ns; 0.01-0.05 = *; 0-001 - 0.01 = **; <0.001 = ***.

### Immunoprecipitation

Cells were transfected in a 100mm dish with 5µg of total DNA with the indicated constructs. After 36h cells were washed two times with 2ml of cold PBS (Biosera, # LM-S20412/500) and lysed in 1 ml of lysis buffer (0.5% NP-40, 1mM EDTA, 150mM NaCl, 50mM Tris, 1mM DTT, 1x Protease Inhibitor Cocktail (Thermo Scientific, #A32961)) for 20min on a rocking platform at 4°C. The lysates were scraped and transferred to a 1.5ml Eppendorf tube, then centrifuged at 14000g for 15min at 4°C. Total protein concentration was measured using Pierce™ BCA® Protein Assay Kits (Thermo Scientific, #23227) and 500µg of proteins were used per IP conditions. From the total lysate 40µl were used as input and mixed with 10µl of 4x Laemmli buffer and stored at -20°C. Anti HA.11-epitope antibody (BioLegend, #901514, 1:150), Cleaved Notch1 (Val1744, Cell Signaling #4147, 1:200), IgG1 control (Thermo Scientific, #A42561, 1:150), and IgG1 rabbit control (Thermo Scientific, #10500C, 1:200) were used for precipitation. Antibody-lysate mixture was incubated at 4°C for 1h in a carousel. Protein G Sepharose™ 4 Fast Flow (Cytiva, #17-0618-01) beads were prepared by washing the resin three times with complete lysis buffer. 50% slurry beads were prepared by mixing equal volumes of resin and lysis buffer. 80µl of beads were added to the antibody-lysate mixture and let them rotate O/N at 4°C. The mixtures were centrifuged at 0.1 RCF, 4°C for 1 min, and the precipitated beads were washed four times with lysis buffer without protease inhibitor. 42µl of 2x Laemmli buffer was added to the beads and boiled at 95°C for 5 min, quickly centrifuged, and analyzed by Western blot.

### Ligand coating plate

96 well plates or 100mm plates were coated with 40µl or 1ml of recombinant Protein G (50µg/ml in PBS) and incubated overnight at RT with gentle rocking movement. Wells were washed three times with PBS at RT and blocked with 40µl or 1ml of BSA (10mg/ml) for 1h at RT. Wells were washed three times with PBS and 40µl or 1ml of recombinant hDLL4-Fc (Biotechne R&D System. #10185-D4) (1µg/ml diluted in 0.1% BSA in PBS) or IgG1-Fc (Thermo Fischer Scientific, # A42561) (1µg/ml diluted in 0.1% BSA in PBS) and incubated for 2h at RT. Coated wells were washed with PBS, and 25.000 cells or 1.75 x 10^6^ were seeded.

### Luciferase assay

Cells were transfected in a 96 well plate with 120ng of total DNA with the indicated constructs after 16h from seeding. After 6h from transfection 150ng/ml of rhWnt3a (R&D Systems, #5036-WN) supplemented media was added to the cell for Wnt activation (L. Shao et al., 2013). 24h after the media was removed and followed by Dual-Glo® Luciferase Assay System (Promega) protocol. Briefly, 40µl of RT complete DMEM was added to the cells for 5 minutes to adapt to the RT. 40 µl of Dual-Glo® Luciferase Reagent was added onto the cells and the whole volume was mixed properly and let them incubate for 25min at RT followed by 5 min of incubation in agitation. Mixture was moved to a white plate (#136101), and the firefly luciferase was measured using Varioscan (#catalog). 40 µl of Dual-Glo® Stop & Glo® Reagent (1/100) was added and mixed well and incubated for 25min at RT followed by 5 min of incubation in agitation. The renilla luciferase luminescence was measured and used to internal normalization. All experiments were conducted, at least, in triplicate, with each experimental value representing the means of three independent replicates. Data were analyzed using GraphPad Prism 11 software. Outliers were identified and removed using the ROUT method (Q = 1%)

### Plasmids

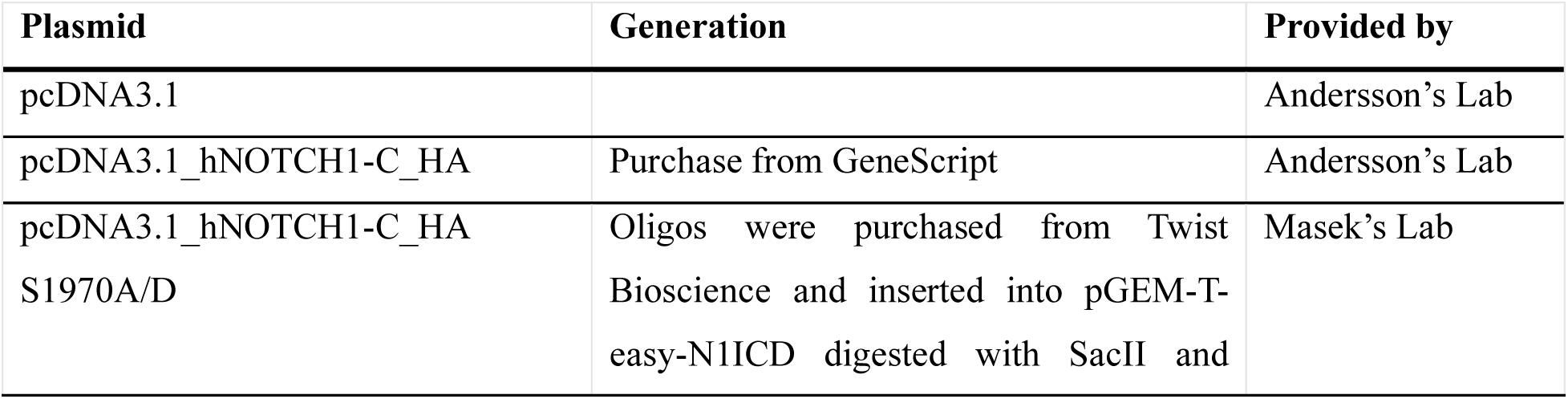
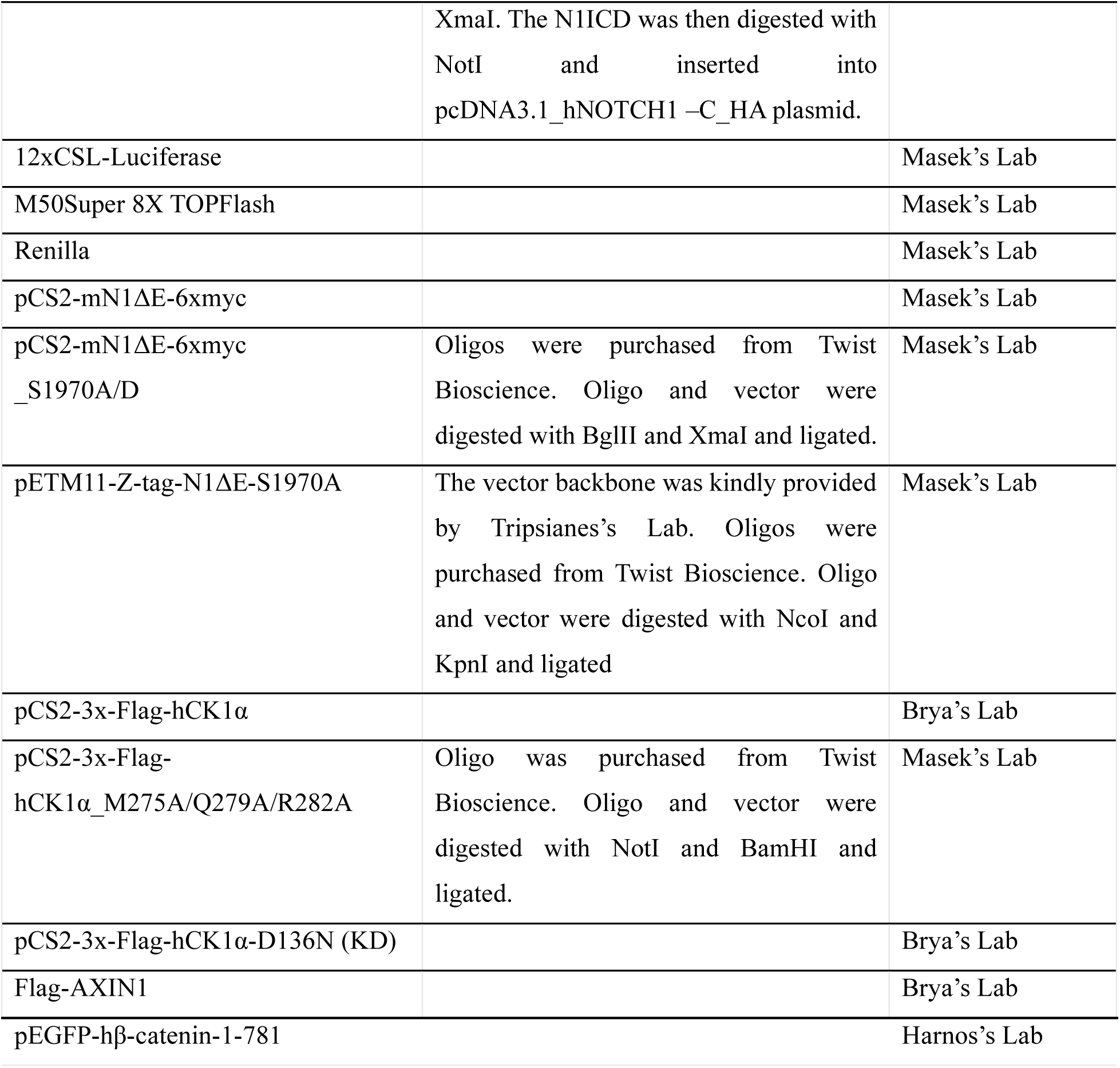

### Transfections

Cells were transfected with the indicated constructs using PEI Prime™ linear polyethylenimine (PEI) 1mg/ml with a 1:3 DNA:PEI ratio. For Luciferase assay each well of the 96-well plate was transfected with total of 120ng of plasmid, including 30ng of 12xCSL-Luc or TOPFlash, 5ng of Renilla, 30ng of N1ΔE or pcDNA3.1, 30ng of hCK1α/hCK1αKD/hAXIN, and topped with pcDNA3.1 empty vector. For Immunoprecipitation, a total of 5µg of plasmid was transfected into a 100mm dish with 2.5µg of the indicated plasmid, topped with pcDNA3.1 empty vector. In a 6-well plate, 500ng of total DNA was transfected with 50ng of the different Wnt components (hCK1α/hAXIN) along with 50ng of N1ΔE and topped with pcDNA3.1 empty vector. For the MG132 and CHX experiments, 500ng of total DNA was transfected with 25ng of N1ΔE. Media was replaced after 6h to minimize the cytotoxicity.

### Western blotting

Cells were washed two times with cold PBS (Biosera, # LM-S20412/500) and lysed in 75-150µl of RIPA buffer (25mM Tris-HCl pH7.6, 150mM NaCl, 1%NP40, 1% Sodium Deoxycholate, 0.1% SDS) for 10min on a rocking platform at 4°C. The lysates were scraped and transferred into a 1.5ml Eppendorf tube and centrifuge at 14000g for 10min at 4°C. Cleared lysate protein concentration was measured using Pierce™ BCA® Protein Assay Kits (Thermo Fisher Scientific, #23227) and 10 to 35µg of proteins were separated by SDS-PAGE (8%) and then transferred to a PVDF (Cytiva #10600021) membrane, using a Trans-Blot Turbo Transfer System (25V, 1.3A, 18 min). Membranes were blocked with TBS-T buffer containing 5% non-fat milk for 1h at RT and then probed with the indicated primary antibody in TBS-T with 1% BSA at 4°C, O/N. Membranes were washed three times for 10min with TBS-T buffer and subsequently incubated with the appropriate secondary antibody (1:20000 in TBS-T buffer with 1% BSA) at RT for 1h. Membranes were then developed with Pierce™ ECL Western Blotting Substrate (Thermo Fisher Scientific, #32209) or SuperSignal™ West Femto Maximum Sensitivity Substrate (Thermo Fisher Scientific, #34096) in a ChemiDoc imager. Western blot images were quantified using ImageLab, with a volumetric approach. β-ACTIN was used for normalization. Briefly, the ratio of POI/β-ACTIN signal intensity was calculated. The Fold change was then calculated using either the control or wild-type condition. Data were then plotted and analyzed using GraphPad Prism 11.

### qPCR

HEK TREx and *CK1α*^-/-^ cells (4 x 10^5^ cells/well) were seeded in 6-well plates. After 24 h, cells were harvested in 1 mL cold TRIzol™ (Thermo Fisher Scientific, #15596026). Total RNA was isolated, and 1µg was treated with DNase I (Thermo Fisher Scientific, #EN0521) for 30 min at 37°C, followed by 1µl of 50mM EDTA for 10 min at 65°C.

cDNA was synthesized using the RevertAid transcriptase (Thermo Fisher Scientific, # EP0442) and subsequently diluted with 80µ of nuclease-free water. qPCR was performed using 2µl of cDNA and Luna® Universal qPCR Master Mix (NEB, #M3003S) on a LightCycler® 480 instrument. Cycling conditions: 42 cycles with a 60°C annealing temperature. Primers used:

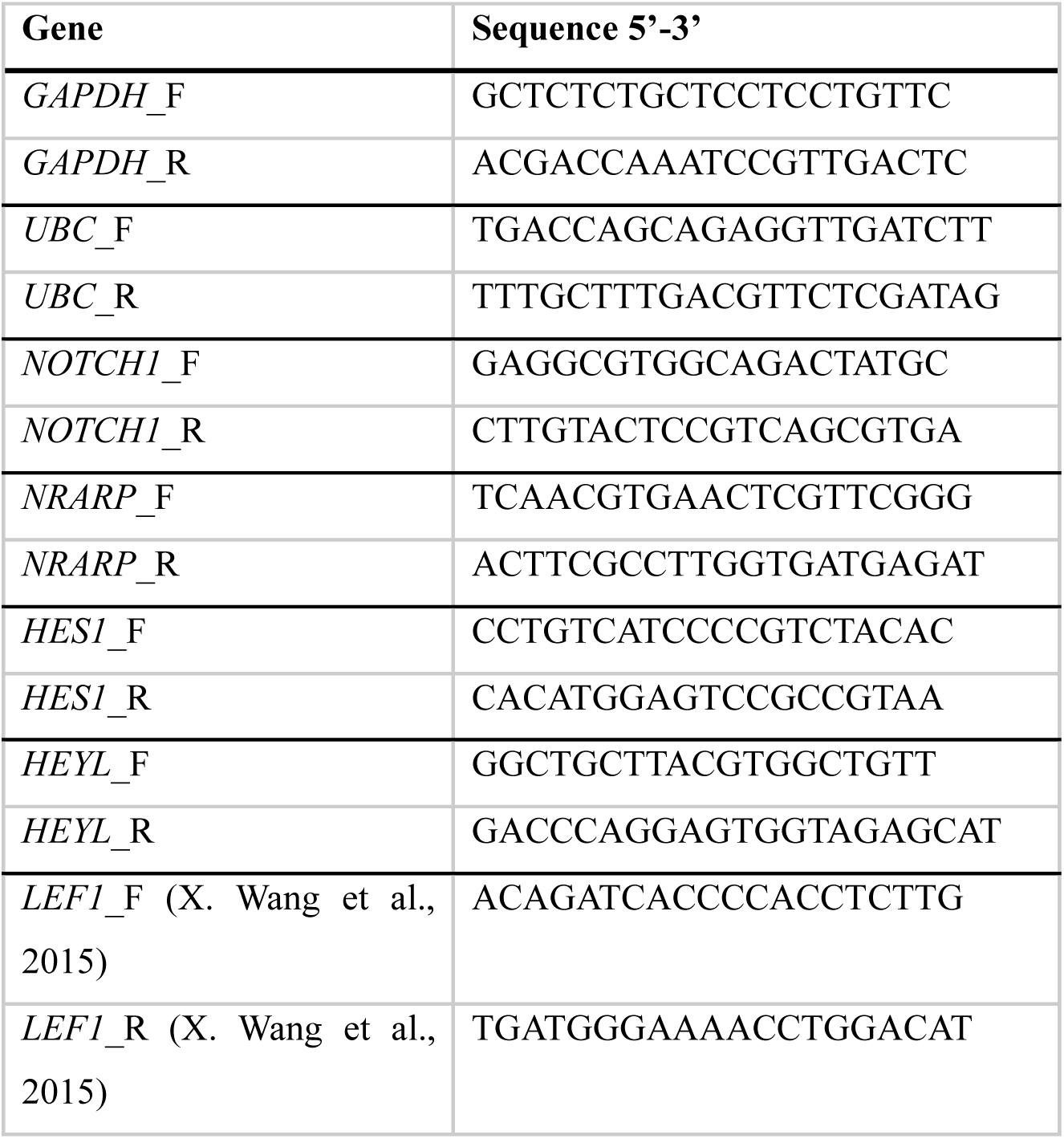

### Protein Digestion

Dry beads from imunnoprecipitation were mixed with 100 mM TEAB (Triethylammonium bicarbonate) in 2% SDC (sodium deoxycholate), 10 mM TCEP (Tris(2-carboxyethyl)phosphine) and 50 mM CAA (chloroacetamide) and shaked for 30 min at 60°C. After incubation IP beads were digested in 50 mM TEAB at 37 °C with 1 µg of trypsin overnight. After digestion, samples were centrifuged and the supernatant was acidified with TFA to 1% final concentration. Supernatant was washed 3times with ethylacetate and the residual ethyacetate was evaporated. Samples were acidified with TFA to 1% final concentration and peptides were desalted using in-house made stage tips packed with C18 disks disks (AttractSPE, Affinisep, France) according to Rappsilber et al (Rappsilber et al., 2007).

### nLC-MS 2 Analysis

Nano Reversed phase column (Ion Opticks, Aurora Ultimate TS 25×75 C18 UHPLC column) was used for LC/MS analysis. Mobile phase buffer A was composed of water and 0.1% formic acid. Mobile phase B was composed of acetonitrile and 0.1% formic acid. Samples were loaded onto the trap column (C18 PepMap100, 5 μm particle size, 300 μm x 5 mm, ThermoScientific) for 4 min at 18 μl/min loading buffer was composed of water, 2% acetonitrile and 0.1% trifluoroacetic acid. Peptides were eluted with Mobile phase B gradient from 4% to 35% B in 60 min. Eluting peptide cations were converted to gas-phase ions by electrospray ionization and analyzed on a Thermo Orbitrap Ascend (Thermo Scientific) by data dependent approach. Survey scans of peptide precursors from 350 to 1400 m/z were performed in orbitrap at 120K resolution (at 200 m/z) with a 100 % ion count target. Tandem MS was performed by isolation at 1,6 Da with the quadrupole, HCD fragmentation with normalized collision energy of 30 % and 10 ms activation time. Fragmentation spectra were acquired in ion trap with scan rate set to Rapid. The MS2 ion count target was set to 150 % and the max injection time was 75 ms. Only those precursors with charge state 2–6 were sampled for MS2. The dynamic exclusion duration was set to 30 s with a 10ppm tolerance around the selected precursor and its isotopes. Monoisotopic precursor selection was turned on. Cycle time was se to 1.5 s.

All data were analyzed and quantified with the MaxQuant software (version 2.4.13.0) (Cox & Mann, 2008). The false discovery rate (FDR) was set to 1% for both proteins and peptides and we specified a minimum peptide length of seven amino acids. The Andromeda search engine was used for the MS/MS spectra search with the Homo sapiens (downloaded from Uniprot in March 2023, containing 20 605 entries). Enzyme specificity was set as C-terminal to Arg and Lys, also allowing cleavage at proline bonds and a maximum of two missed cleavages. Carbamidomethylation of cysteine was selected as fixed modification and N-terminal protein acetylation and methionine oxidation as variable modifications. Data analysis was performed performed using Perseus 1.6.15.0 software (Tyanova et al., 2016) and GO terms analysis using RStudio.

Intersection cluster analysis

Unique proteins were identified using the filtering >2 value per condition rule. GO analysis of single and double clusters was done using Rstudio

### Recombinant protein expression and purification

Recombinant N1ICD-ANK wild-type (WT) and S1970A mutant proteins (amino acids 1872-2126) were expressed from pETM11 vectors as fusion proteins containing an N-terminal 6×His tag followed by a Z-tag and a tobacco etch virus (TEV) protease recognition site (ENLYFQG). Recombinant CK1ε (catalytic core, amino acids 1-301) was expressed from pETM11 vector with a TEV-cleavable N-terminal 6×His tag. Plasmids were transformed into chemically competent E. coli BL21(DE3) cells by heat shock. Bacterial cultures were grown under antibiotic selection (30 µg/mL kanamycin for N1ICD-ANK; 30 µg/mL kanamycin for CK1ε) in M9 minimal medium supplemented with 0.5 g/L ^15^NH_4_Cl (CN80P10, Cortecnet) as the sole nitrogen source (for N1ICD-ANK) or in LB medium (for CK1ε). Cells were grown at 37 °C to an OD of 0.7, shifted to 16 °C for 1 h, and expression was induced with 0.5 mM isopropyl β-D-1-thiogalactopyranoside (IPTG; A1008, PanReac AppliChem), followed by overnight incubation at 16 °C. Cells were harvested by centrifugation (4,000 × g, 15 min, 4 °C) and resuspended in lysis buffer (for N1ICD-ANK: 50 mM Tris-HCl pH 8.0, 150 mM NaCl, 10 mM imidazole, 20% w/v sucrose, 5 mM DTT; for CK1ε: 50 mM Tris-HCl pH 7.2, 500 mM NaCl, 10 mM imidazole, 10% v/v glycerol, 300 mM sucrose, 1% NP-40, 3 mM β-mercaptoethanol) supplemented with complete EDTA-free Protease Inhibitor Cocktail (11873580001, Merck) and phenylmethanesulfonyl fluoride (PMSF; 329-98-6, AppliChem). Cell suspensions were stored at -20 °C. After thawing, suspensions were lysed by sonication and clarified by centrifugation (27,000 × g, 40 min, 4 °C). For N1ICD-ANK purification, clarified lysate was applied to a NiNTA column (HiTrap IMAC HP 5 mL, Cytiva) equilibrated with binding buffer (50 mM Tris-HCl pH 8.0, 150 mM NaCl, 10 mM imidazole). Protein was eluted with a 10-500 mM imidazole gradient. The N-terminal tag was cleaved by incubation with TEV protease (purified in-house) for 1 h at room temperature, followed by overnight dialysis at 4 °C against binding buffer. The sample was reapplied to a Ni-NTA column, and the tag-free protein was collected in the flow-through and concentrated to a final volume of 1 mL using a centrifugal concentrator (Vivaspin 20, 10,000 MWCO, Sartorius). Final purification was performed by size exclusion chromatography on a Superdex 75 Increase 10/300 GL column (Cytiva) equilibrated with NMR buffer (50 mM sodium phosphate pH 6.5, 50 mM KCl, 1 mM DTT). Protein concentration was determined by absorbance at 280 nm using an extinction coefficient of 11,460 M -1 cm -1 (identical for WT and S1970A). After addition of 1 mM d_6_-EDTA, protein aliquots were flash-frozen and stored at -80 °C. For CK1ε purification, clarified lysate was applied to a Ni-NTA column equilibrated with binding buffer (50 mM Tris-HCl pH 7.2, 500 mM NaCl, 10 mM imidazole, 10% v/v glycerol, 3 mM β-mercaptoethanol). Protein was eluted with a 10-500 mM imidazole gradient. The N-terminal tag was cleaved by TEV protease during overnight dialysis against cation-exchange binding buffer (50 mM Tris-HCl pH 7.2, 50 mM NaCl, 5 mM EDTA, 5% v/v glycerol, 3 mM β-mercaptoethanol). The dialyzed sample was applied to a cation exchange column (HiTrap SP 5 ml, Cytiva). Protein was eluted with a 50-1000 mM NaCl gradient and concentrated to a final volume of 1 mL. Final purification was performed using a Superdex 75 Increase 10/300 GL column equilibrated with NMR buffer. Protein concentration was determined using an extinction coefficient of 43,320 M -1 cm -1.

### NMR spectroscopy

NMR measurements were performed on 850 MHz Bruker Avance NEO spectrometer equipped with ^1^H/ ^13^C/^15^N TCI cryogenic probe with z-axis gradients. For real-time NMR monitoring of N1ICD-ANK phosphorylation, 100 µM of ^15^N-labelled N1ICD-ANK WT or S1970A was mixed with 2 µM CK1ε in NMR buffer (50 mM sodium phosphate pH 6.5, 50 mM KCl, 1 mM DTT) supplemented with 1 mM d_6_ -EDTA, 2 mM ATP, 10 mM MgCl_2_, and 10% D_2_O (for the deuterium lock). Phosphorylation was initiated by addition of CK1ε directly into the NMR tube, and ^1^H-^15^N HSQC spectra were acquired every 30 minutes for 14 h at 25 °C. Spectra were processed using TopSpin 4.0.6 and analysed in NMRFAM-SPARKY (Lee et al., 2015) and in-house Python scripts.

### Liquid chromatography-tandem mass spectrometry (LC-MS/MS)

Proteins were digested with trypsin in 50 mM ammonium bicarbonate at 40 °C for 2 h. The peptide mixtures were analyzed via liquid chromatography-tandem mass spectrometry (LC-MS/MS) using an Ultimate 3000 RSLCnano system (Thermo Fisher Scientific, Waltham, MA, USA) coupled to an Impact II Qq-Time-of-Flight mass spectrometer (Bruker, Bremen, Germany). Peptides were separated on an Acclaim Pepmap100 C18 column (3 µm particle size, 75 μm × 500 mm; Thermo Fisher Scientific) using the following solvent gradient (mobile phase A: 0.1% FA in water; mobile phase B: 0.1% FA in 80% ACN; 300 nL/min flow rate): elution started with 1% mobile phase B, gradually increasing to 56% over 50 min, then increasing linearly to 80% mobile phase B over the next 5 min, followed by isocratic elution with 80% mobile phase B for 10 min. The outlet of the analytical column was connected directly to the CaptiveSpray nanoBooster ion source (Bruker, Bremen, Germany). MS and MS/MS spectra were acquired using a data-dependent strategy with a 3-s cycle time. The mass range was set to m/z 150–2200, with precursor ions selected from m/z 300–2000. MS and MS/MS acquisition speeds were 2 Hz and 4–16 Hz, respectively, with MS/MS acquisition rate varying based on precursor intensity. Mass spectrometric data preprocessing, including recalibration, compound detection, and charge deconvolution, was performed using DataAnalysis software (version 4.2 SR1; Bruker).

Mascot MS/MS ion searches (Matrixscience, London, UK; version 2.5.1) were done against in-house database containing expected protein sequences. cRAP contaminant database (downloaded from http://www.thegpm.org/crap/) was also searched to exclude contaminant spectra. Mass tolerances for precursor and MS/MS fragment ions were 10 ppm and 0.1 Da, respectively, with the option of one 13C atom to be present in the parent ion. Oxidation of methionine and phosphorylation of serine, threonine and tyrosine were set as variable modifications; no fixed modifications were set. The modification degree was determined from peak areas in extracted ion chromatograms of tryptic peptide forms covering the individual amino acids in the protein sequence.

### Molecular dynamics simulations

Two simulation systems were prepared from a trimeric protein complex derived from pdb:3V79 (DNA deleted): one with the native serine at position 1970 of chain K (non-phosphorylated), and one with a phosphoserine (SEP) substitution at the same position (phosphorylated). For the phosphorylated system, residue SER197 of chain K was manually renamed to SEP in the PDB file and the phosphate group atoms (P, O1P, O2P, O3P) were verified to match the naming convention required by the phosaa14SB parameter set. Both systems were prepared using the LEaP module of AmberTools. The AMBER14SB force field (leaprc.protein.ff14SB) was used for standard amino acids, supplemented with the phosaa14SB parameter set for the phosphoserine residue, derived from Dries et al. (DOI: 10.1021/acs.jctc.4c00732). Each system was solvated in a truncated octahedral box of TIP3P water molecules with a minimum solute-to-box-edge distance of 8 Å. The net system charge was neutralised with Na+ or Cl− counterions as appropriate, after which additional NaCl ion pairs were added to approximate a physiological ionic strength of 0.15 M. Topology (.prmtop) and coordinate (.inpcrd) files were saved for use in OpenMM.

Simulation Protocol

All molecular dynamics simulations were performed using OpenMM 8.4. AMBER topology and coordinate files were loaded directly via AmberPrmtopFile and AmberInpcrdFile, with box vectors set from the inpcrd file to correctly initialise the periodic boundary conditions. Long-range electrostatics were treated with Particle Mesh Ewald (PME) using a real-space cutoff of 1.0 nm and an Ewald error tolerance of 5 × 10−4. Bonds involving hydrogen atoms were constrained using the HBonds scheme, and water molecules were treated as rigid bodies, permitting a 2 fs integration timestep. Each system was energy-minimised for up to 1,000 steps using the L-BFGS algorithm implemented in OpenMM. Following minimisation, the system was equilibrated in the NVT ensemble for 10 ps with velocities initialised from a Maxwell-Boltzmann distribution at 300 K, using a Langevin middle integrator with a friction coefficient of 1 ps−1. Constant pressure was then introduced by adding a Monte Carlo barostat targeting 1 bar, with volume moves attempted every 25 steps. NPT equilibration was then continued for a further 50 ps at 300 K and 1 bar. Production simulations were run in the NPT ensemble at 300 K and 1 bar for 500 ns each. Trajectory coordinates were saved every 10 ps (50,000 steps) in DCD format. Thermodynamic state variables (potential energy, kinetic energy, temperature, volume, density) were recorded at the same interval. System checkpoints were saved every 100 ps. Simulations were performed on an NVIDIA L40S GPU using mixed-precision arithmetic.

### *Xenopus* experiments

All procedures involving *Xenopus laevis* were conducted in accordance with Czech legislation on the use of animals for research and were approved by the relevant institutional and governmental authorities (MSMT-21426/2025, Ministry of Education, Youth and Sports of the Czech Republic; 45980/2023-MZE-13143, Ministry of Agriculture of the Czech Republic; and MZP/2025/630/2482, Ministry of the Environment of the Czech Republic).

#### Embryo handling

Embryos were obtained and maintained using standard procedures. Briefly, testes were surgically removed from anesthetized adult males (20% MS-222; Sigma-Aldrich, A5040) and stored in cold 1× Marc’s Modified Ringer’s solution (MMR; 100 mM NaCl, 2 mM KCl, 1 mM MgSO₄, 2 mM CaCl₂, 5 mM HEPES, pH 7.4) supplemented with 50 µg/mL gentamicin (Sigma-Aldrich, G3632). To induce ovulation, sexually mature females were injected with 260 U of human chorionic gonadotropin (hCG; Merck, Ovitrelle 250G) into the dorsal lymph sac and kept overnight at 18 °C. The following day, eggs were collected by gently squeezing the females and fertilized in vitro using a freshly macerated piece of testis in 0.1× MMR. Fertilized embryos were cultured in 0.1× MMR at 18–21 °C and staged according to Zahn et al., 2022

#### mRNA synthesis and microinjection

Capped synthetic mRNAs, encoding NICD WT, NICD S1970A, and NICD S1970D (all sourced from Masek lab), together with CK1α (Gybeľ et al., 2024), and β-catenin (source from Harnos lab), were generated by in vitro transcription using linearized plasmid templates and a commercial mRNA transcription kit (mMessage mMachine T7 and SP6) according to the manufacturer’s instructions (Thermo Fisher Scientific). mRNA quality and concentration were verified by agarose gel electrophoresis and spectrophotometry.

#### Secondary axis assay

Embryo microinjections were performed at the 4-cell stage in 3% Ficoll 400 (Cytiva, #17-0300-10) prepared in 0.5× MMR. Approximately 10 nl of mRNA solution (containing 250 pg of β-catenin, NICD WT, NICD A and NICD D; 25 pg of NICD and CK1α) was injected per blastomere using a calibrated microinjection system. For the secondary axis assay, mRNA was injected into equatorial region of both ventral blastomeres, as indicated in the figure legends. After injection, embryos were cultured in 0.1× MMR until the desired stages. For analysis of secondary axis formation, embryos were allowed to develop until late neurula stages (NF stage 19), fixed in 4% formaldehyde in PBS (Sigma-Aldrich), and scored under the Leica S9i stereomicroscope. Embryos were categorized as having no secondary axis, partial secondary axis, or complete secondary axis. Representative images were captured using a standard bright-field imaging setup, using Zeiss AxioZoom.V16 microscope.

#### Explant preparation and gene expression analysis

For transcriptional analysis, NICD and/or CK1α mRNAs were injected into animal pole of all four blastomeres at the 4-cell stage. Embryos were cultured until blastula (NF stage 8), and animal cap explants were dissected using Dumont #5 Fine Forceps (11254-20) in 0.5× MMR supplemented with 50 µg/mL gentamicin (Sigma-Aldrich, G3632). Explants were cultured further until NF stage 10.5. Total RNA was isolated using the RNA isolation kit (CathGene - MR20250). First strand cDNA was synthesized using RevertAid Reverse Transcriptase (Thermo Fisher), from 0.5 mg of total RNA. Semiquantitative RT-PCR was performed using PCRBIO Taq DNA Polymerase (PCR Biosystems). Gene-specific primers were used to assess expression of the Notch target gene Hes4 (hairy2 in *Xenopus laevis*). ef1α was used as a cDNA loading control. Primer sequences for hairy2 and ef1α were obtained from here (Agius et al., 2000; Murato et al., 2007). PCR amplification was performed using 28 and 25 cycles, respectively. For each experimental condition of one given experiment, 10-15 animal caps were pooled and analyzed. Non-injected sibling embryos serve as a positive control in the first lane of PCR gel. Expression levels were normalized to the housekeeping gene ef1α and analyzed using the GraphPad Prism, Tukey’s multiple comparisons test. All experiments were performed in four independent biological replicates. Data are presented as mean with SD. Statistical significance was determined using appropriate parametric tests as indicated in the figure legends, with p < 0.05 considered significant.

## Author Contribution

How each author was involved with the manuscript (e.g., ^1^study concept and design; ^2^acquisition of data; ^3^ analysis and interpretation of data; ^4^drafting of the manuscript; ^5^critical revision of the manuscript for important intellectual content; ^6^statistical analysis; ^7^obtained funding; ^8^technical, or material support; ^9^study supervision) ^123456789^

**FT**^1234567^, **LMA**^23456^, **HH**^23456^, **JS**^23^, **WD**^23^, **MD**^1,3^, **GC**^1458^, **TG**^58^, **KP**, **ERA**^8^, **VB**^58^, **PP**^358^, **KT**, **JH**^134579^, **JM**^1345679^

## ACKNOWLEDGMENTS

We are grateful to Marian Novotny, Faculty of Science, Charles University for introducing us to the AlphaFold modeling and interpretation of the results. LCMS analyses were performed in Laboratory of Mass Spectrometry at Biocev research center, Faculty of Science, Charles University. Confocal miscroscopy was performed with help from Viničná Microscopy Core Facility; Faculty of Science, Charles University. AlphaFold Metacentrum was used for modelling prediction https://metavo.metacentrum.cz/en/myaccount/pubs. CIISB, Instruct-CZ Centre of Instruct-ERIC EU consortium, funded by MEYS CR infrastructure project LM2023042 and European Regional Development Fund-Project „Innovation of Czech Infrastructure for Integrative Structural Biology“ (No. CZ.02.01.01/00/23_015/0008175), is gratefully acknowledged for the financial support of the measurements at the CEITEC Proteomics Core Facility. Computational resources were provided by the e-INFRA CZ project (ID:90254), supported by MEYS CR

