## Supplementary figures for "NOTCH1-specific phosphorylation of S1970 by Casein Kinase 1 is required for NOTCH1 transcriptional competence and signaling activity *in vivo*"

**Author Contribution**: How each author was involved with the manuscript (e.g., [1]study concept and design; [2]acquisition of data; [3] analysis and interpretation of data; [4]drafting of the manuscript; [5]critical revision of the manuscript for important intellectual content; [6]statistical analysis; [7]obtained funding; [8]technical, or material support; [9]study supervision) [123456789]

**FT**[1234567], **LMA**[23456], **HH**[23456], **JS**[23], **WD**[23], **MD**[1,3], **GC**[1458], **TG**[58], **KP**, **ERA**[8], **VB**[58], **PP**[358], **KT**, **JH**[134579], **JM**[1345679]

### SUPPLEMENTARY DATA:

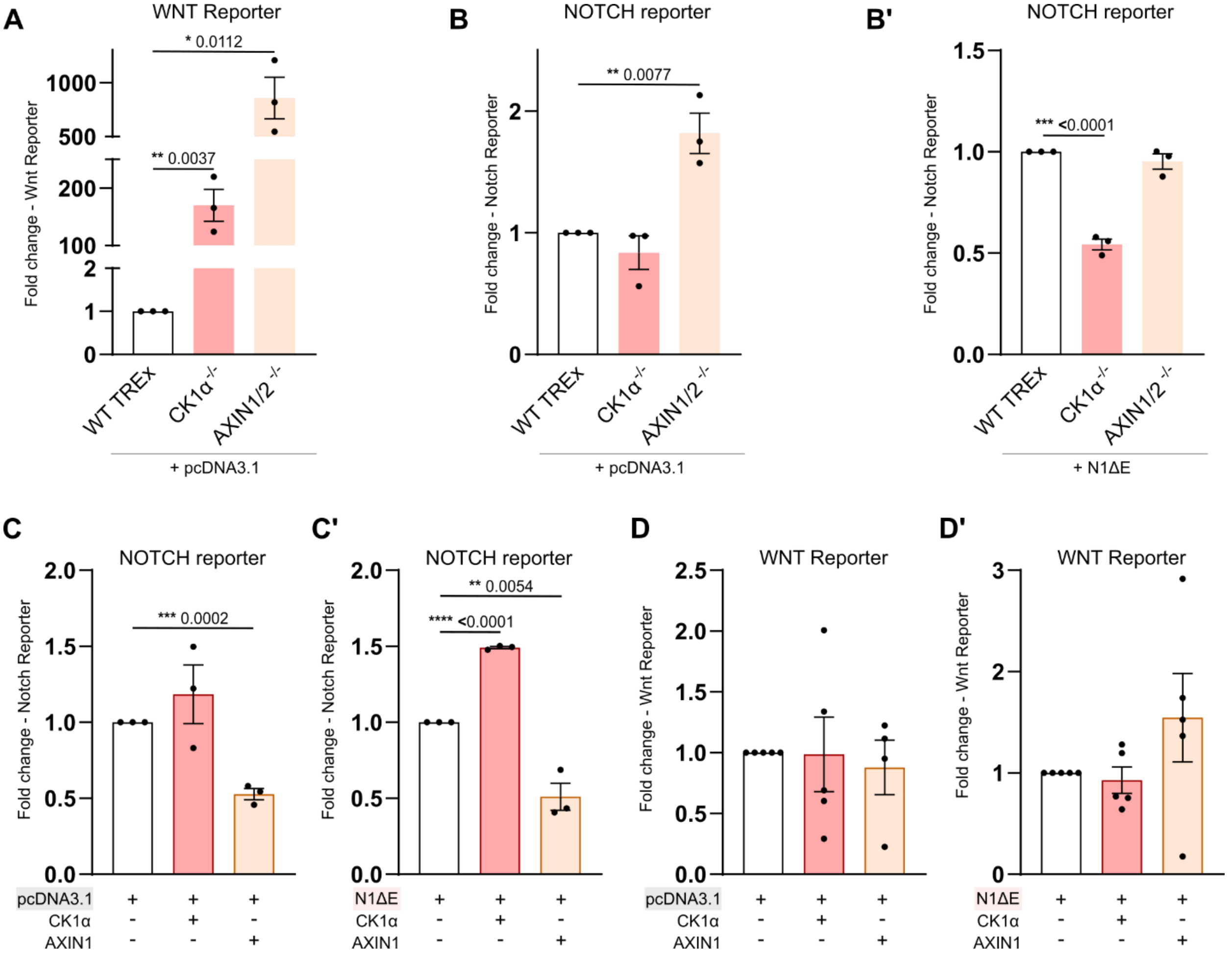


***Supplementary Figure 1: CK1α associates with the Notch to sustain signaling and regulate NICD stability*** ***(A)*** *WNT luciferase reporter assay in* $CK1\alpha^{-/-}$ *cells transfected with pcDNA3.1.* ***(B-B')*** *Luciferase Notch reporter assay in* $CK1\alpha^{-/-}$ *cells transfected with either pcDNA3.1 or NIΔE plasmids.* ***(C-C')*** *Luciferase Notch reporter assay in HEK293T cells transfected with either CK1α or AXIN1 and either pcDNA3.1 or NIΔE.* ***(D-D')*** *Luciferase Wnt reporter assay in HEK293T cells transfected with either CK1α or AXIN1 and either pcDNA3.1 or NIΔE. Each dot represents a biological replicate (mean of a technical triplicate). Unpaired-T test (** $P \leq 0.05$; ** $P \leq 0.01$; *** $P \leq 0.001$, **** $P > 0.0001$*, ns = not significant) Bars represent SEM.*

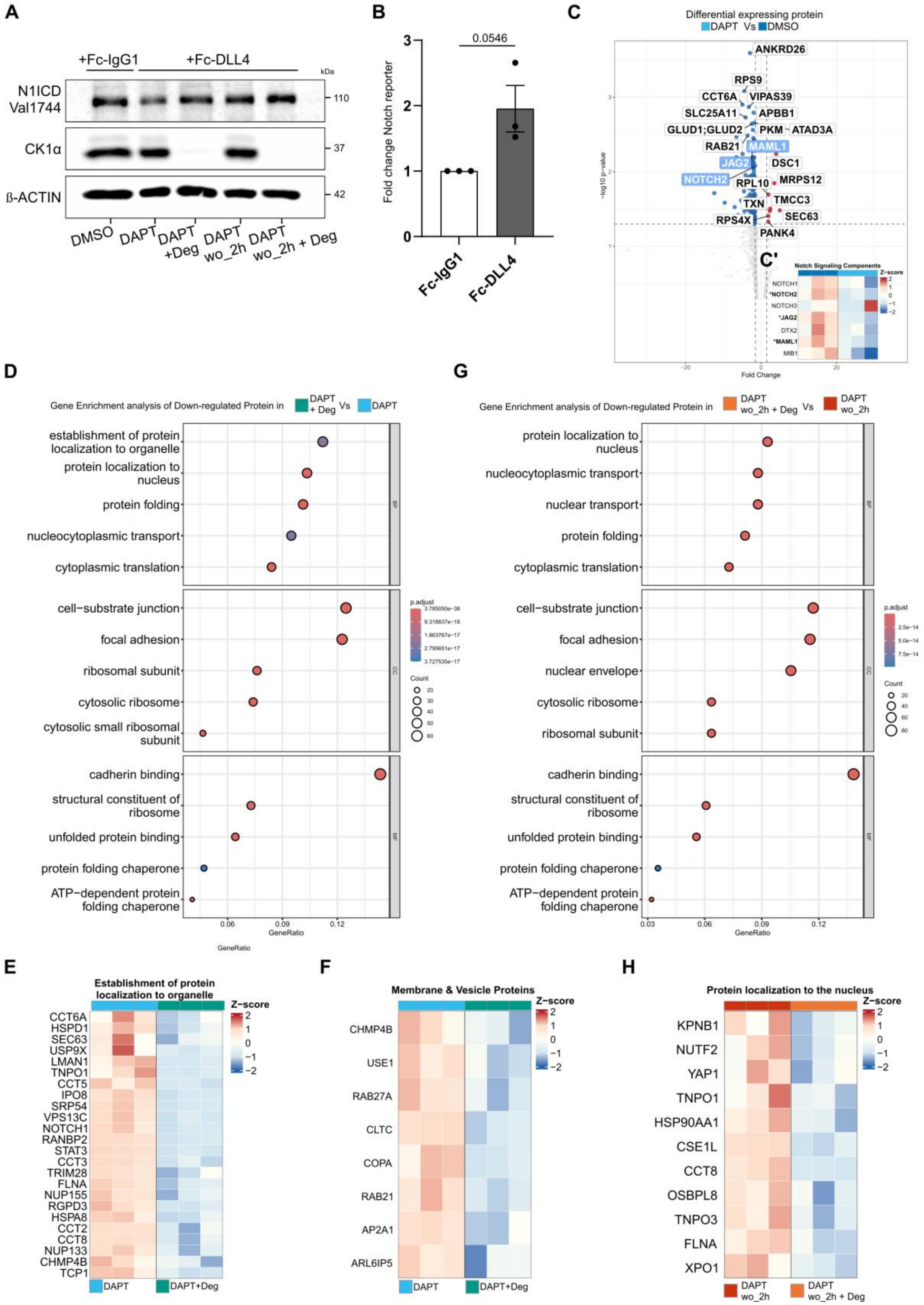
A
+Fc-IgG1
+Fc-DLL4
kDa
N1ICD Val1744
CK1α
β-ACTIN
110
37
42
DMSO
DAPT
DAPT +Deg
DAPT wo_2h
DAPT wo_2h + Deg
B
0.0546
Fold change Notch reporter
Fc-IgG1
Fc-DLL4
C
Differential expressing protein
DAPT Vs DMSO
ANKRD26
RPS9
CCT6A
VIPAS39
SLC25A11
APBB1
GLUD1;GLUD2
PKM
ATAD3A
RAB21
MAML1
JAG2
DSC1
NOTCH2
RPL10
MRPS12
TXN
TMCC3
RPS4X
SEC63
PANK4
-log10 p-value
Fold Change
C'
Notch Signaling Components
Z-score
NOTCH1
*NOTCH2
NOTCH3
*JAG2
DTX2
*MAML1
MIB1
D
Gene Enrichment analysis of Down-regulated Protein in DAPT + Deg Vs DAPT
establishment of protein localization to organelle
protein localization to nucleus
protein folding
nucleocytoplasmic transport
cytoplasmic translation
cell−substrate junction
focal adhesion
ribosomal subunit
cytosolic ribosome
cytosolic small ribosomal subunit
cadherin binding
structural constituent of ribosome
unfolded protein binding
protein folding chaperone
ATP−dependent protein folding chaperone
p.adjust
Count
GeneRatio
G
Gene Enrichment analysis of Down-regulated Protein in DAPT wo_2h + Deg Vs DAPT wo_2h
protein localization to nucleus
nucleocytoplasmic transport
nuclear transport
protein folding
cytoplasmic translation
cell−substrate junction
focal adhesion
nuclear envelope
cytosolic ribosome
ribosomal subunit
cadherin binding
structural constituent of ribosome
unfolded protein binding
protein folding chaperone
ATP−dependent protein folding chaperone
p.adjust
Count
GeneRatio
E
Establishment of protein localization to organelle
Z−score
CCT6A
HSPD1
SEC63
USP9X
LMAN1
TNPO1
CCT5
IPO8
SRP54
VPS13C
NOTCH1
RANBP2
STAT3
CCT3
TRIM28
FLNA
NUP155
RGPD3
HSPA8
CCT2
CCT8
NUP133
CHMP4B
TCP1
DAPT
DAPT+Deg
F
Membrane & Vesicle Proteins
Z−score
CHMP4B
USE1
RAB27A
CLTC
COPA
RAB21
AP2A1
ARL6IP5
DAPT
DAPT+Deg
H
Protein localization to the nucleus
Z−score
KPNB1
NUTF2
YAP1
TNPO1
HSP90AA1
CSE1L
CCT8
OSBPL8
TNPO3
FLNA
XPO1
DAPT wo_2h
DAPT wo_2h + Deg

***Supplementary Figure 2: CK1α degradation impairs Notch trafficking proximity***
*(**A**) Western blot of the Cleaved NOTCH1 ICD (Val1744) and CK1α in hNOTCH1-UltraID cell line treated as in Fig. 3D. (**B**) Luciferase Notch reporter assay in hNOTCH1-UltraID cells seeded in Fc-IgG1 control or Fc-DLL4 plate. (**C**) Volcano plot visualization of the DEP in DAPT wo_2h Vs DAPT groups, and (**C'**) heatmap visualization of the Notch signaling components identified. (**D**) Dot plot visualization of Gene Ontology (BP, CC, MF) of the top 5 down-regulating proteins in DAPT + CK1α-Deg Vs DAPT alone. (**E**) Heatmap visualization of Non membrane bounded organelle assembly and (**F**) Membrane & Vesicle proteins. (**G**) Dot plot visualization of Gene Ontology (BP, CC, MF) of the top 5 down-regulating proteins in DAPT wo_2h + CK1α-Deg Vs DAPT wo_2h alone. (**H**) Heatmap visualization of Protein localization to the nucleus proteins. Volcano plot represents significant DEPs Log2 Fold Change $\geq$1.5, -log10 p-value $\geq$ 1.3. Each dot represents a biological replicate; Unpaired-T test; Bars represent SEM*

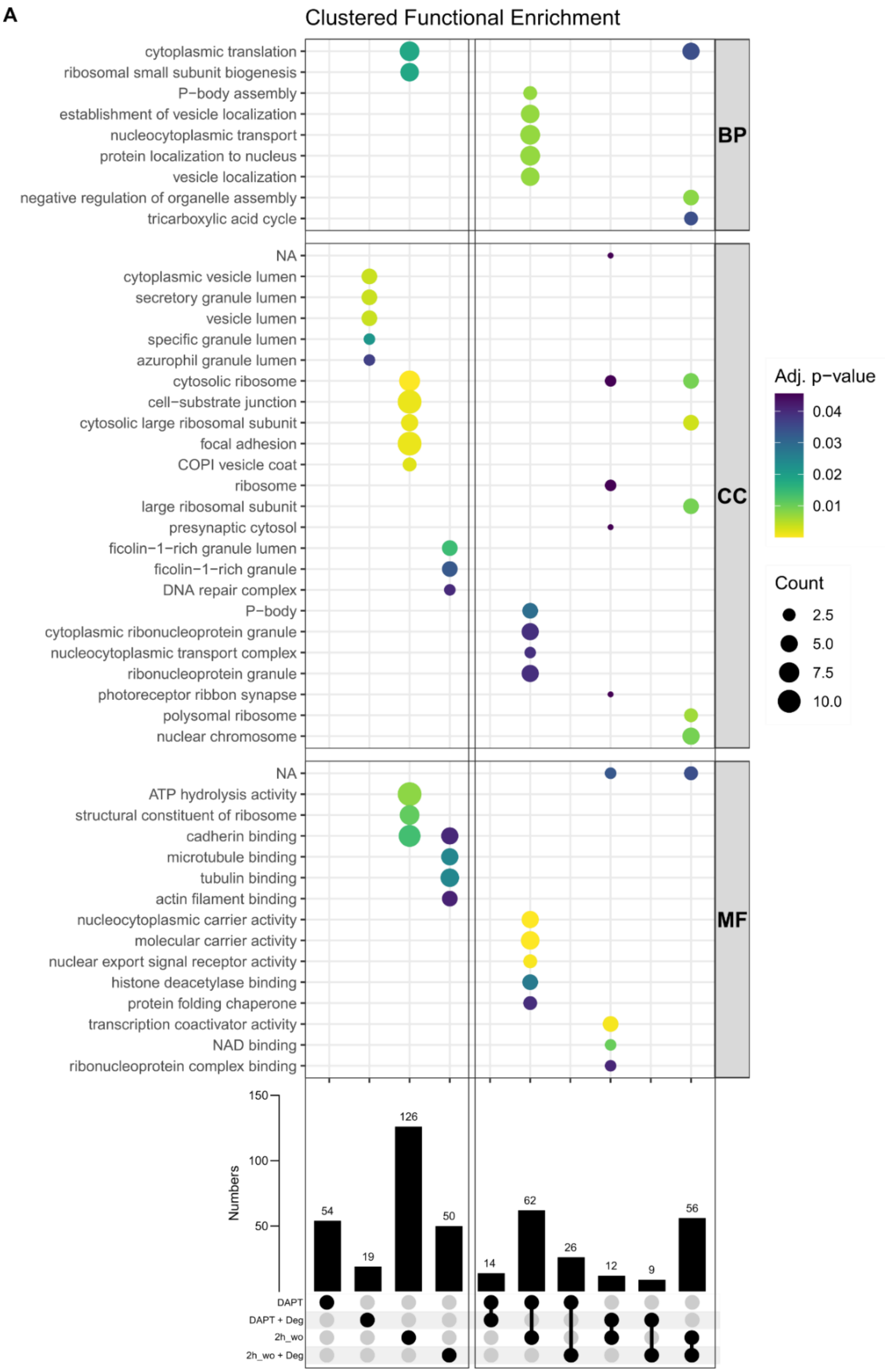


***Supplementary Figure 3: Analysis of proteins identified in single and intersected cluster***

*(A) Dot plot visualization of Gene Ontology (BP, CC, MF) analysis of proteins identified in single or intersected cluster groups represented as a black dot (lower panel).*

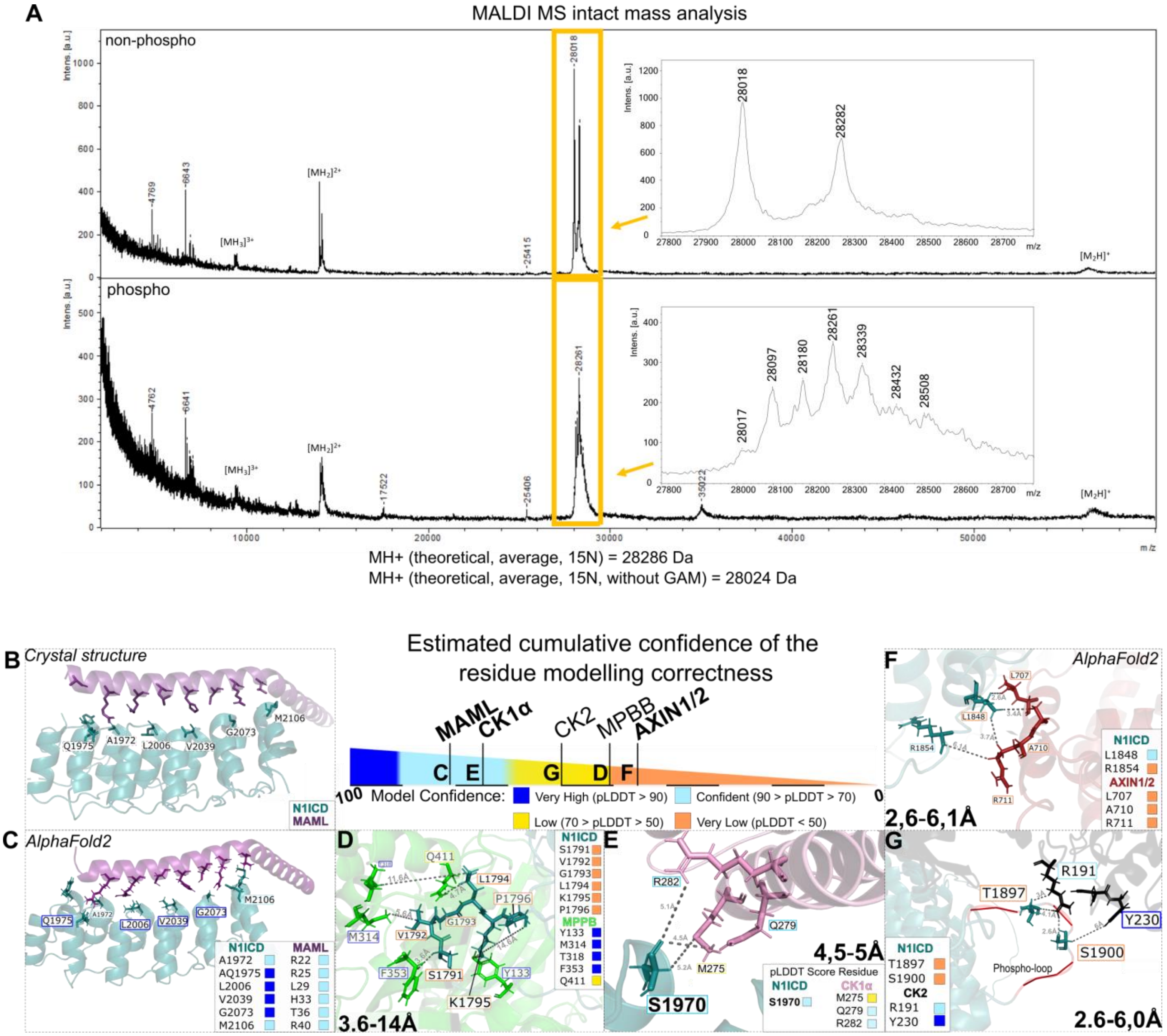


***Supplementary Figure 4: MALDI MS and AlphaFold 2.0 modelling prediction and validation*** ***(A)*** *MALDI MS intact mass analysis of N1ICD ANK core alone (non-phospho, top), or after phosphorylation with CK1 core (phospho, bottom).* ***(B)*** *Crystal structure (PDB 3V79) from Choi et al., 2012. and (**C**) AlphaFold modelling of N1ICD and MAML1 interaction. (**D-G**) AlphaFold modelling prediction interaction of N1ICD with (D) MPPB, (E) CK1α, (F) AXIN1/2, and (G) CK2 proteins. pLDDT, predicted local distance difference test.*

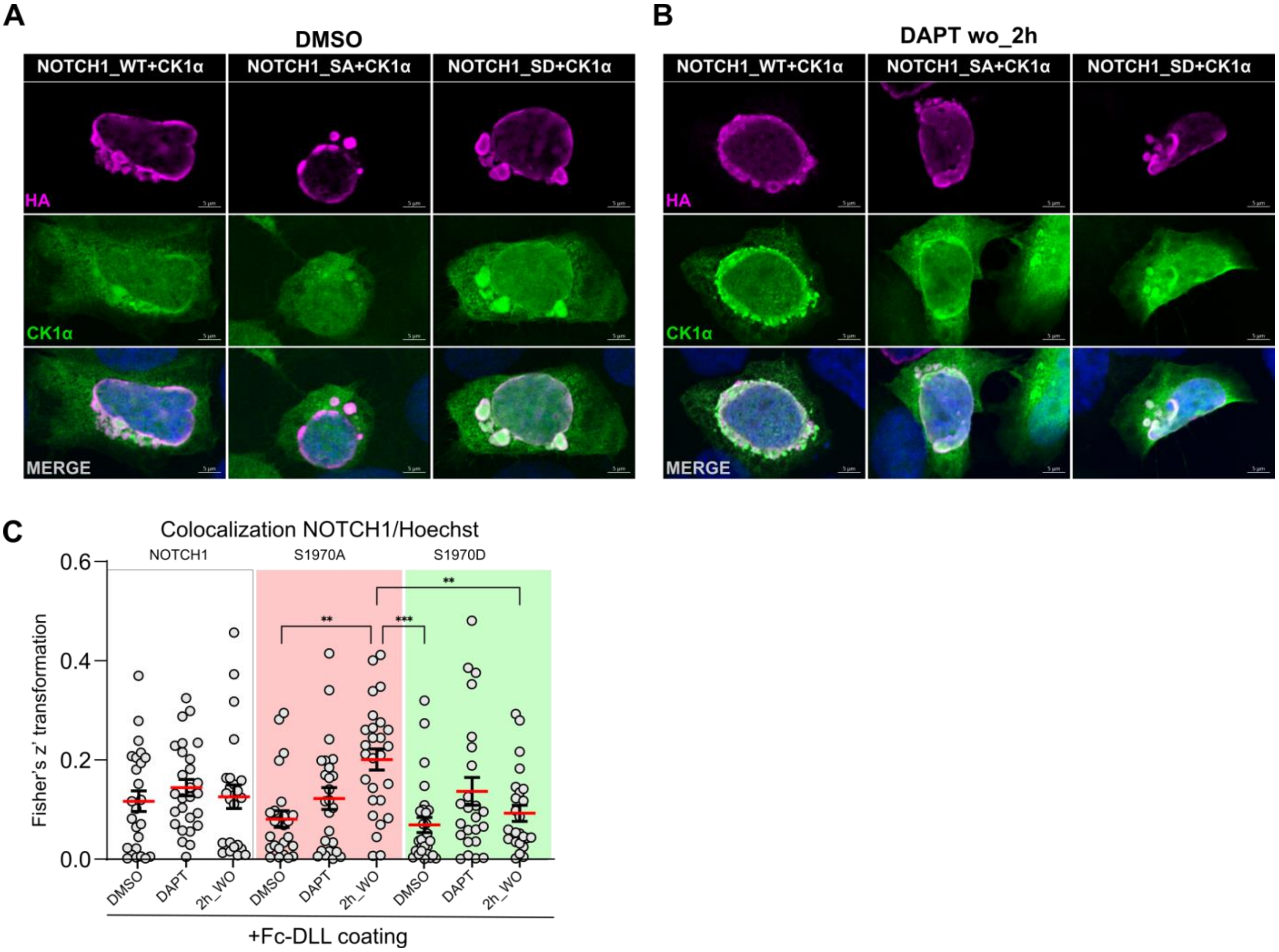


***Supplementary Figure 5: S1970A mutation influences NOTCH1 nuclear localization after DAPT washout***

*(**A-B**) Representative confocal images (MIP) of DMSO (A) and DAPT wo_2h-treated (B) cells transfected with NOTCH1-S1970A/D HA and 3x-FlagCK1α. (**C**) Fisher's z' quantification of NOTCH1 (WT, S1970A, S1970D) and Hoechst (nuclei) colocalization in N1-4KO cells seeded on a Fc-DLL4 coated plates. Scale bars: 5µm. Ordinary One-way ANOVA; ***P < 0.001, ****P < 0.0001.*

**A**

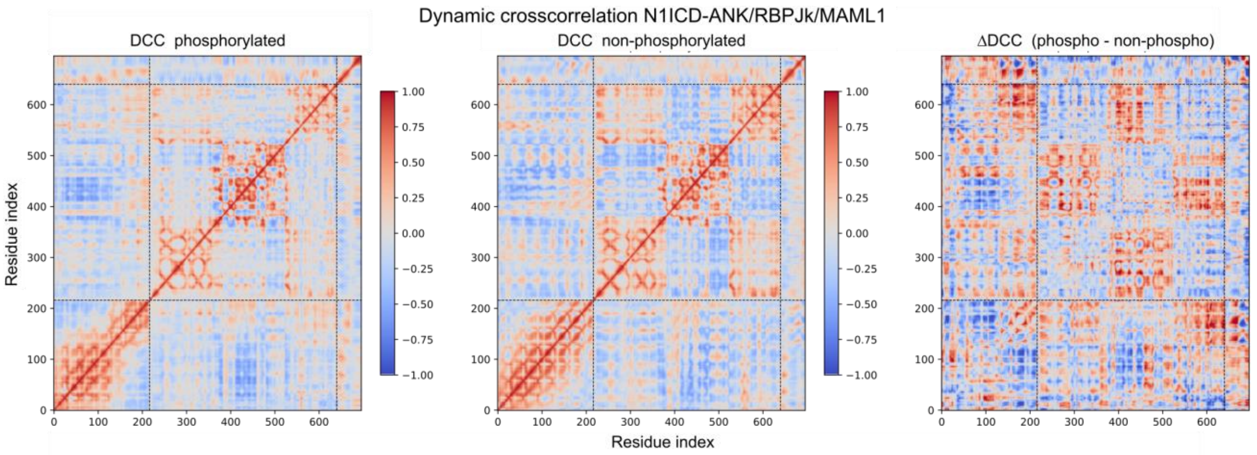


***Supplementary Figure 6: Dynamic cross-correlation analysis of N1ICD-ANK S1970 phosphorylation reveals a shift in ternary complex association***

*(**A**) Dynamic cross-correlation (DCC) analysis of phosphorylation-induced shifts.*
